# Intensive task-switching training and single-task training differentially affect behavioral and neural manifestations of cognitive control in children

**DOI:** 10.1101/2023.12.22.573065

**Authors:** Sina A. Schwarze, Corinna Laube, Neda Khosravani, Ulman Lindenberger, Silvia A. Bunge, Yana Fandakova

## Abstract

The ability to flexibly switch between tasks develops during childhood. Children’s task-switching performance improves with practice, but the underlying processes remain unclear. We used functional magnetic resonance imaging to examine how nine weeks of task-switching training affect performance and task-related activation and functional connectivity. Children (8–11 years) were assigned to one of three groups: intensive task switching (SW; n = 72), intensive single tasking (SI; n = 74), and passive control (n = 41). While mixing costs decreased in both training groups initially, only the SW group maintained these training-related improvements at the end of training. Activation in dorsolateral prefrontal cortex decreased with training, but again only the SW group maintained these activation decreases at the end of training. Condition-specific connectivity increases with task switching became less pronounced with training, especially in the SI group. Lower costs of task switching along with decreased task-related activations suggest increased processing efficiency in frontoparietal regions with training. Intensive task-switching training was associated with sustained changes, possibly facilitated by a greater mismatch between processing supplies and environmental demands. Our findings suggest that experience-dependent changes with intensive task-switching training do not mirror maturational processes but rather facilitate performance via more efficient task processing.

## 1. Introduction

Executive functions describe a set of control processes supporting goal-directed behavior (Diamond 2013). Task switching, the ability to flexibly switch between different tasks, constitutes a key component of executive functions (Miyake et al. 2000; Miyake and Friedman 2012) and continues to improve across childhood (Cepeda et al. 2001; Crone et al. 2004; Reimers and Maylor 2005; Crone, Bunge, et al. 2006; Huizinga and van der Molen 2007; Weeda et al. 2014). Accordingly, a number of studies have aimed to improve children’s task-switching abilities with training (Karbach and Kray 2009; Kray, Karbach, Haenig, et al. 2012; Zinke et al. 2012; Kray et al. 2013; Dörrenbächer et al. 2014; Karbach et al. 2017; Zuber et al. 2023). These studies have consistently demonstrated that performance in the trained tasks improves with practice. To date, the mechanisms underlying these training-related task-switching improvements in childhood are not well understood. Specifically, it is unclear whether cognitive training has a similar impact on the neural processes associated with task switching as age-dependent maturation (cf. Jolles and Crone 2012) or whether different, potentially child-specific neural processes underlie training-related improvements in task switching.

The present study sought to disentangle these hypotheses by examining the sequential progression of training-related changes in the neural processes underlying task switching. We focused on children between 8 and 11 years because executive functions continue to develop during this age period (Reimers and Maylor 2005; Karbach and Unger 2014; see also Tervo-Clemmens et al. 2023) and may therefore be particularly malleable through training (Luna et al. 2015; Laube et al. 2020). Combining neuroimaging methods with intensive task-switching training can help elucidate the dynamics of neural change with experience, thereby extending our understanding of task-switching and executive functioning development in childhood.

To examine task switching in the laboratory, task-switching paradigms require individuals to perform two or more tasks in an intermixed fashion, such that each trial constitutes either a repeat of the previous task or a switch to a different one (cf. Koch and Kiesel 2022). The demand to switch to a different task elicits performance costs (i.e., switch costs), evident in lower accuracy and longer response times (RTs). Switch costs are assumed to reflect the updating of the relevant task set and the inhibition of the no-longer relevant task set (e.g., Allport et al. 1994; Rogers and Monsell 1995; Meiran 1996; Mayr and Kliegl 2000; Wylie and Allport 2000). Blocks of trials involving task switches can further be compared to single blocks in which participants perform the different tasks separately. The comparison of repeat trials in mixed blocks to single-block trials (i.e., mixing costs) captures the processes supporting the intermixed performance of tasks, in particular the maintenance and monitoring of multiple task sets (e.g., Rubin and Meiran 2005; Pettigrew and Martin 2016).

Compared to young adults, children show both greater mixing and switch costs. However, while switch costs approach adult levels around age 10 years, age differences in mixing costs continue to be evident up to adolescence (Cepeda et al. 2001; Crone et al. 2004; Reimers and Maylor 2005; Crone, Bunge, et al. 2006; Huizinga and van der Molen 2007; Manzi et al. 2011; Schwarze et al. 2023). This pattern suggests that the neural processes supporting transient task-set updating and inhibition mature earlier than the processes supporting the sustained maintenance and monitoring of multiple task sets (Kray et al. 2004; Schwarze et al. 2024).

### 1.1 Task-switching training

Task-switching performance can be improved with training, at least over the short term, across the lifespan (Kray and Lindenberger 2000; Cepeda et al. 2001; Minear and Shah 2008; Berryhill and Hughes 2009; Strobach et al. 2012; von Bastian and Oberauer 2013; Dörrenbächer et al. 2014). During task-switching training, participants typically practice switching between tasks over the course of several sessions. Training studies have consistently revealed improved task-switching performance on the trained tasks across various participant groups (Kray and Dörrenbächer 2020). By comparing performance improvements with task-switching training to an active control group that practices the same tasks in a single-task condition, studies have further demonstrated that it is specifically switching between tasks that leads to improved performance, as opposed to repeated practice with the task rules in separate blocks (Minear and Shah 2008).

In studies with adults, mixing costs were substantially reduced or even eliminated upon training (Berryhill and Hughes 2009; Strobach et al. 2012), while switch costs were mostly reduced but remained present after training (Kray and Lindenberger 2000; Cepeda et al. 2001; Strobach et al. 2012). This pattern of results suggests that training may be particularly effective for reducing the demands on task-set maintenance and monitoring processes associated with mixing costs (Rubin and Meiran 2005; Pettigrew and Martin 2016). From a developmental perspective, these results stress the potential of training to mitigate age differences in task switching, which are particularly pronounced with respect to mixing costs (Cepeda et al. 2001; Reimers and Maylor 2005).

Indeed, task-switching training in children leads to improvements in both mixing and switch costs (Cepeda et al. 2001; Karbach and Kray 2009; Kray, Karbach, Haenig, et al. 2012; Zinke et al. 2012; Kray et al. 2013; Dörrenbächer et al. 2014; Karbach et al. 2017; Zuber et al. 2023). Some studies showed greater training-related reductions in mixing costs in children than in adults (Cepeda et al. 2001; Karbach and Kray 2009; Karbach et al. 2017). These findings of larger training benefits in the task-switching component that demonstrates protracted development suggest that task-switching skills may be especially malleable through training in middle childhood, when the corresponding neural processes have not yet fully matured (cf. Wass et al. 2012; Kühn and Lindenberger 2016).

While previous studies demonstrated that training reduced mixing and switch costs in children, facilitating more adult-like levels of task-switching performance, no studies to date have examined the neural basis of these experience-dependent changes. It is therefore unclear whether neural activation associated with task switching also becomes more adult-like with training or whether children improve their performance through different mechanisms.

### 1.2 Neural changes associated with training

Task switching has been associated with increased activation in frontoparietal brain regions (for recent meta-analyses, see Worringer et al. 2019; Zhang et al. 2021), in particular in the inferior frontal junction (IFJ; cf. Derrfuss et al. 2005), the superior parietal lobe (SPL), and the dorsolateral prefrontal cortex (dlPFC), along with functional connections among them (Yin et al. 2015; Dajani et al. 2020). Training studies of task switching or dual tasking in adults have revealed that activation in these regions decreases with training (Dux et al. 2009; Jimura et al. 2014; Garner and Dux 2015), consistent with training studies of executive functions in general, which also showed decreases in brain activation with training (Landau et al. 2004; Landau et al. 2007; Dux et al. 2009; Schneiders et al. 2011; Jimura et al. 2014). At the same time, other executive-function training studies have reported training-related increases in activation (Olesen et al. 2004; Erickson et al. 2007; Westerberg and Klingberg 2007; Jolles et al. 2010; Schweizer et al. 2013; Buschkuehl et al. 2014).

As a result, the overall picture of training-induced quantitative changes in activation in adults is mixed (Landau et al. 2004; Kelly and Garavan 2005; Buschkuehl et al. 2012; Hsu et al. 2014; Constantinidis and Klingberg 2016). Training-related decreases in frontoparietal activation have been interpreted as improved efficiency of rule processing, in the form of more precise and faster recruitment of the neural circuit the task relies on, sparser neural representations, or both (Poldrack 2000; Kelly and Garavan 2005; Kelly et al. 2006). During task switching, such faster and more differentiated rule processing would be especially important for performance in mixed blocks in which multiple task sets must be simultaneously and flexibly managed, resulting in reduced mixing costs. In contrast, training-related activation increases have been interpreted as stronger involvement of the brain regions that support effective task execution (Poldrack 2000; Kelly and Garavan 2005; Kelly et al. 2006). Similarly, functional connectivity among frontoparietal regions has been found to increase with cognitive training, both at rest (Jolles et al. 2013; Mackey et al. 2013; Guerra-Carrillo et al. 2014) and during task performance (Kundu et al. 2013; Thompson et al. 2016). Connectivity increases might allow for faster reconfiguration of task sets in prefrontal regions (cf. Richter and Yeung 2014). Building on these investigations in adults, the present study sought to shed light on the neural processes supporting training-induced improvements in task switching in children aged between 8 and 11 years.

Based on the existent cognitive training literature, we hypothesized two alternative patterns of neural change with training in children. On one hand, children might show reduced activation in frontoparietal brain regions, similar to what has been observed in training studies with adults (Landau et al. 2004; Landau et al. 2007; Dux et al. 2009; Schneiders et al. 2011; Jimura et al. 2014). Training-related reductions in frontoparietal activation have been previously demonstrated with attention training in children: using electroencephalography, Rueda et al. (2012) showed faster recruitment of the attention network after training in 4–6-year-olds.

On the other hand, cognitive training in children may have similar effects on neural processes as age-dependent maturation (Jolles and Crone 2012), such that with training, children’s brain activation becomes increasingly similar to the activation seen in adults (Rueda et al. 2005; Jolles et al. 2012). Consistent with this hypothesis, studies have reported more adult-like connectivity patterns following a working-memory training in children (Astle et al. 2015; but see Jolles et al. 2013) or after a combined executive-function training in adolescents (Lee et al. 2022).

During task switching, children recruit similar brain regions as adults, but they modulate task-related activation with increasing switching demands less adaptively (Bunge and Wright 2007; Velanova et al. 2008; Wendelken et al. 2012; Mogadam et al. 2018; Engelhardt et al. 2019; Kupis et al. 2021; Zhang et al. 2021; Schwarze et al. 2023; but see Crone, Donohue, et al. 2006; Morton et al. 2009). For example, in the same sample of children prior to training, Schwarze and colleagues (2023) demonstrated that children showed a smaller upregulation of activation than adults in frontoparietal regions during mixed compared to single blocks, and during switch compared to repeat trials (see also Wendelken et al. 2012; Crone, Donohue, et al. 2006; Schwarze et al. 2024). Thus, if we observed more pronounced increases of frontoparietal activation with greater task-switching demands after training, this pattern would be consistent with previously observed patterns of age-dependent maturation.

Finally, in addition to these quantitative changes, children may also show qualitative changes by recruiting additional or different brain regions to meet increased demands on task switching during training (Buschkuehl et al. 2012; Jolles and Crone 2012). This would speak to fundamentally different training effects in children compared to adults, potentially due to the continuing development of the underlying neural circuitry (Galván 2010).

### 1.3 Present study

In the present study, we examined the sequential progression of training-related neural changes in two groups of 8–11-year-olds who trained with different amounts of task switching over nine weeks. To elucidate trajectories of change beyond pre- and post-training measures (cf. Lindenberger and Lövdén 2019; Lövdén et al. 2020), participants performed task switching in the magnetic resonance imaging (MRI) scanner or MRI simulator on two occasions during the training period in addition to the pre- and post-training sessions. We predicted that task-switching performance would improve with training in both groups, albeit at different time points during training. Specifically, consistent with previous findings, we predicted that intensive task-switching training would induce performance improvements relatively early on during training. Training the task rules during predominantly single tasking might also improve performance early in training, albeit to a smaller extent (Karbach and Kray 2009). As the training duration of the present study was longer than in previous studies, we hypothesized that it might allow children performing mainly single-task training to catch up, later in training, with their peers who mainly trained task switching. Given children’s presumed difficulty in binding rules with appropriate actions (Karbach and Kray 2009), the present design allowed us to investigate whether a regime that allows for establishing these bindings through more intensive single-task practice may be as (or even more) beneficial as an intensive practice of switching between tasks.

We further sought to test the hypotheses regarding neural changes in task-based activation and connectivity outlined above. On one hand, greater increases in activation for higher mixing and switch demands after training would speak to training evoking similar changes as age-dependent maturation. Conversely, training-related decreases in frontoparietal activation would suggest more efficient rule processing in these regions, consistent with previous training studies with adults. We further expected increasing connectivity among frontoparietal regions with training, along with potentially smaller differences between conditions. Mirroring our hypotheses of training-related changes in performance, training-related change in activation and connectivity might be more extensive or become evident earlier in children practicing intensive task-switching compared to children practicing intensive single-tasking. Finally, given that mixing costs and the associated neural processes show larger age differences in the examined age group, we expected that training-related neural changes would be more pronounced for mixing than for switch costs.

## 2. Materials and Methods

The hypotheses and plans for analysis were preregistered before the start of data analysis at https://osf.io/by4zq/.

### 2.1 Research participants and study overview

A total of 187 children aged between 8 and 11 years (M = 9.96 years, SD = 0.70, 90 girls, 97 boys) were pseudo-randomly assigned to one of three groups: two training groups and a passive control group. An overview of the study design is depicted in Figure 1A.

**Figure 1.**
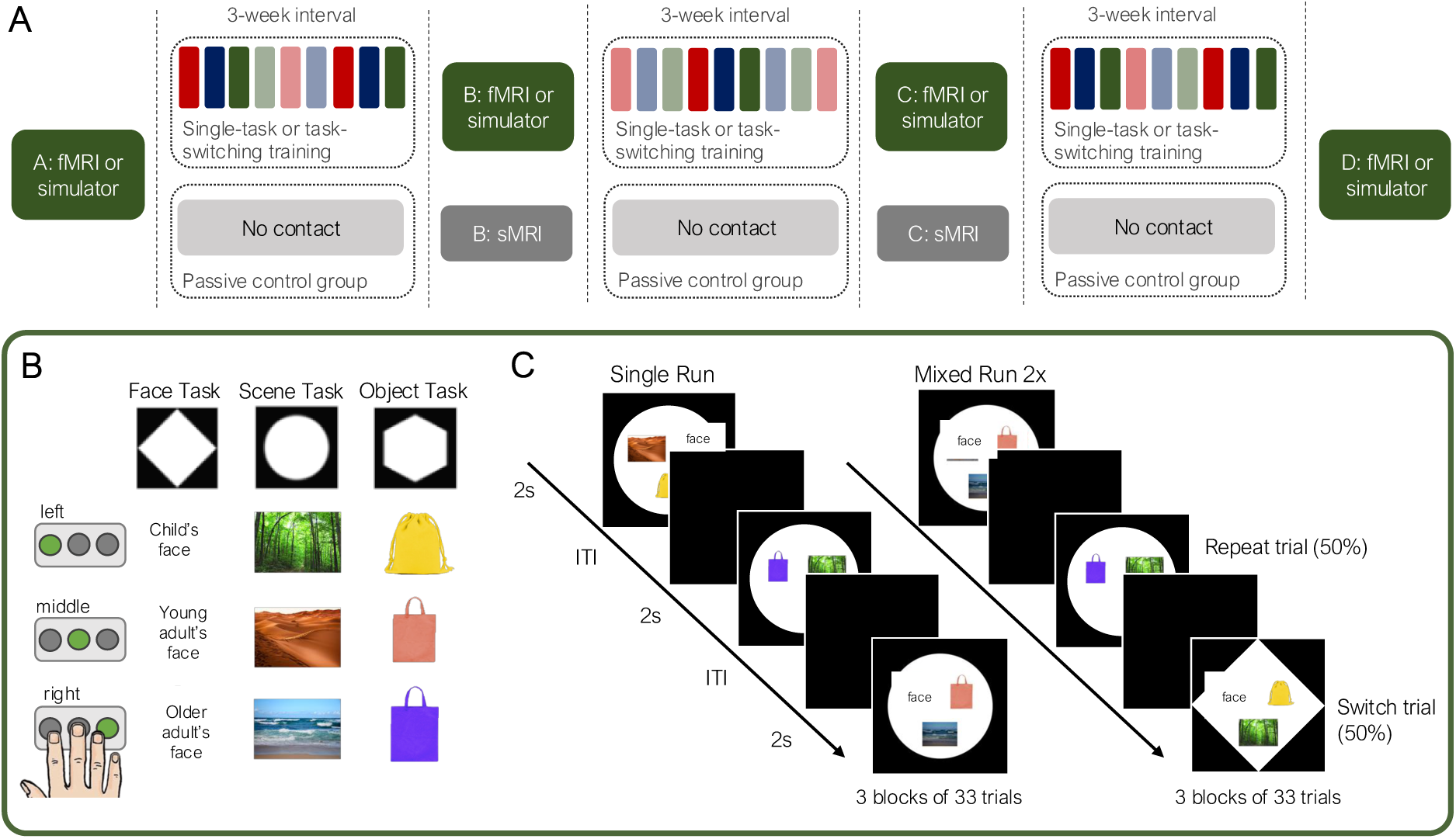
Outline of study design and experimental task-switching paradigm. (A) The timeline of training and assessment across the nine weeks for the three groups. fMRI or simulator indicates that the main task-switching paradigm (see B and C) was performed in the MRI scanner or MRI simulator, sMRI indicates structural MRI scans only. The colored bars indicate the training games: opaque colored bars indicate one of the three repeating games and translucent bars indicate one of the unique games, with the color indicating which of the repeating tasks matched the structure of the unique task. (B) The task-switching paradigm that all groups completed in the fMRI scanner or simulator. The shape cue indicated one of the three tasks. As indicated by three exemplar stimuli of each task, participants selected one of three buttons based on the face’s age in the Face Task, the type of environment in the Scene Task and the color of the object in the Object Task. (C) Showing three sequential trials of the single and mixed blocks; in the single block depicted here, participants performed the scene task on every trial. In the mixed task, the shape cues (and therefore tasks) repeated on some trials and switched on others. ITI: inter-trial interval. Image credits: Young and old adult faces were taken from the FACES collection (Ebner et al. 2010). B & C: Adapted from <u>Schwarze et al. (2023)</u>, Figure 1, under <u>CC.BY 4.0</u>.

The two training groups practiced for nine weeks on a tablet at home for a total of 27 training sessions (30–40 min per session). In each session, participants completed a task-switching training game. An intensive task-switching (SW) training group (N = 72, 36 girls, 36 boys; age: M = 9.85 years, SD = 0.64) completed 17% single-task blocks and 83% mixed-task blocks per training game. An intensive single-tasking (SI) training group (N = 74, 34 girls, 40 boys; age: M = 9.86 years, SD = 0.70) trained on 83% single-task blocks and 17% mixed-task blocks per training game. The stimuli and rules in each game were identical between the two training groups such that the groups differed only in the relative demands on task switching (see section 2.2). The passive control (PC) group (N = 41, 20 girls, 21 boys; age: M = 10.30 years, SD = 0.72) did not perform any training games. Children in the PC group were slightly older than in the SI group (est. = –0.48; 95%-CI = –0.75, –0.21) and the SW group (est. = –0.50; 95%-CI = –0.77, –0.23), who did not differ from each other (est. = –0.01; 95%-CI = –0.23, 0.20).

All participants spoke German as their primary language and had no history of psychological or neurological diseases. All participants who provided MRI data were right-handed and were screened for MRI suitability. Parents and children provided informed written consent. All participants were reimbursed with 10€ per hour spent at the laboratory. The training groups received an additional bonus of 40€ for the completion of all training games and MRI/MRI-simulator sessions. Additionally, children in the training groups received a toy as a reward for their performance on the training games (see details below). The study was approved by the ethics committee of the Freie Universität Berlin and conducted in line with the Declaration of Helsinki.

In addition to at-home training, both training groups performed four sessions of a task-switching paradigm (described in section 2.3) in the MRI scanner or MRI simulator: before training (pre-test, session A), after approximately three weeks of training (session B), after approximately six weeks of training (session C), and again after approximately nine weeks, when the training was completed (post-test, session D). No MRI data were collected for the MRI simulator participants, who performed the task-switching paradigm in a mock scanner that looked just like the MRI scanner. The PC group performed the same task-switching paradigm in the MRI scanner at sessions A and D, while sessions B and C only included structural scans.

Behavioral analyses were based on the four (for the SW and SI group) or two (for the PC group) sessions of the experimental task-switching paradigm performed in the MRI scanner or simulator. To ensure that participants included in the analyses performed this paradigm meaningfully, we excluded data in a session-specific manner based on preregistered performance criteria. Specifically, for each session, if a child performed below 50% accuracy in the run of single blocks (run 1, see below for more details on the paradigm) or below 35% accuracy in either of the two runs of mixed blocks (run 2 and 3), their data from that session were excluded from analyses. Additionally, we excluded 4 participants (2 from each training group) from all analyses because they did not complete at least half of the 27 training games. Based on these criteria, behavioral analyses included 162 children at session A (SW = 61 [10 excluded based on session-specific performance], SI = 65 [8], PC = 36 [5]), 117 at session B (SW = 58 [9 excluded], SI = 59 [9]), 116 at session C (SW = 55 [7 excluded], SI = 61 [8]), and 135 at session D (SW = 49 [14 excluded], SI = 57 [10]), PC = 29 [7]).

Of the children included in behavioral analyses, we additionally excluded children from neuroimaging analyses based on in-scanner head motion. Functional MRI (fMRI) volumes with framewise displacement (Power et al. 2012) above 0.4 mm were labeled as low-quality (cf. Dosenbach et al. 2017). If any of the fMRI runs of a specific session exceeded 50% of low-quality volumes, the session was excluded for that participant. Thus, fMRI analyses included 87 children at session A (SW = 33 [3 excluded], SI = 30 [9], PC = 24 [7]), 56 at session B (SW = 33 [2 excluded], SI = 23 [8]), 55 at session C (SW = 31 [4 excluded], SI = 24 [6]), and 71 at session D (SW = 23 [5 excluded], SI = 24 [5]), PC = 24 [1]).

Note that two of the four participants who were excluded from all analyses because they had completed too few training games were included in the neuroimaging analysis for session A, on which we defined the regions of interest (ROIs). Twenty-one participants left the study after session A (SW = 6, SI = 6, PC = 9).

### 2.2 At-home training

Children in the SW and SI groups received a tablet after their first MRI or MRI simulator session and were instructed to complete three training games per week for nine weeks (i.e., 27 games in total; Figure 1A; see Supplementary Table 1 for details of training games). The training games on the tablet were programmed using Unity (Version 5.6.1; Unity Technologies). Completion of the games was self-paced; however, a new game only became available 24 hours after the completion of the previous game. Three games were repeated every other week in the same order (i.e., five repetitions of each of the three games across training). One of the repeating games was identical to the paradigm performed in the MRI or MRI-simulator sessions. Each repetition of the three games was interspersed with three unique games that were performed only once (for a total of 12 unique games). The unique games were designed to have the same rule structure as one of the repeating games, while using different stimuli. Each game started with task instructions followed by 3 practice blocks of 15 trials each, during which feedback was provided. No feedback was given during the rest of the game.

Two thirds (i.e., 18) of the training games consisted of 486 trials, one third (i.e., 9) consisted of 485 trials, resulting in a minimally different number of single trials performed at each game (see Supplementary Table 2 for details). In each game, the SW group completed 17% single-block trials and 83% mixed-block trials, while the SI group completed 83% single-block trials and 17% mixed-block trials. For both groups, mixed blocks included 50% repeat and switch trials with unpredictable cues that appeared simultaneously with the target. For all games, each trial lasted up to 3 s and responses had to be given within this period, with stimuli presentation ending when a response was given. There was a 50 ms interval between response and presentation of the next trial. After each block, children could decide independently when to start the next block by pressing a button. Each game lasted between 30 and 40 minutes.

To encourage the completion of the games, children received stars at the end of each block that were converted into coins at the end of each game. Children could trade the coins for toys at any of the MRI or MRI-simulator sessions, with a greater number of coins allowing children to receive larger toys. The number of stars received after each block depended on accuracy, with bonus points being awarded for faster responses compared to the previous blocks as long as performance did not drop below 80% accuracy for the SW group and 90% accuracy for the SI group. On average, children in the SW and SI group completed 25.2 (SD = 3.52) and 25.4 (SD = 3.02) training games, respectively (no difference between groups: t = 0.00, p = 1). Additionally, the two training groups did not differ in the overall number of coins they earned over the course of training (t = 0.39, p = .70; mean number of coins: SW = 366.04, SI = 359.84). Thus, children in the two training groups completed similar amounts of training and earned similar rewards.

### 2.3 Experimental task-switching paradigm

For the training groups, all four laboratory sessions included a task-switching paradigm that participants performed in the MRI scanner or MRI simulator (see Schwarze et al. 2023, for detailed paradigm description). Participants were familiar with the paradigm from an assessment session completed prior to the first MRI or MRI-simulator session, and two practice blocks completed in the MRI simulator right before the actual task.

The task-switching paradigm consisted of three tasks: the Face Task, the Scene Task, and the Object Task. Participants had to perform the task cued by the shape of the background, based on previously learned rules linking each shape with one of the three tasks (Figure 1B). Specifically, the Face Task required the presented face to be categorized by age (child, young adult, or older adult; two different images for each age group), the Scene Task required the presented scene to be categorized by its location (forest, desert, or ocean; two different images for each location), and the Object Task required the presented object to be categorized by color (yellow, red, or purple; two different images for each color). Responses were given via button press with three fingers of the right hand. The stimuli and the task cue appeared at the same time. The three cues, the 18 stimuli (i.e., six per task), and the mapping between stimuli and response buttons were identical at all MRI sessions. The arrangement of the target images varied randomly on each trial independent of the categorization rule.

In each session, participants performed 3 runs of 99 trials each (Figure 1C). Every trial lasted 2 s, followed by a fixation cross (1–6s, jittered) along with an extended fixation period (20 s) after every 33 trials. Participants were instructed to respond as quickly and accurately as possible. Responses that were given during stimulus presentation and within the first second of the ITI were included in the analyses (see below). In the first run (i.e., single run), tasks were presented sequentially in a single-task manner. In runs 2 and 3 (i.e., mixed runs), the three tasks were intermixed with a switch rate of 50%, and switches were unpredictable. The first trial of each run was excluded from all analyses. The sequence in which the tasks were presented in the single run differed between sessions. Similarly, the pseudo-random order of task presentation in the mixed runs differed between sessions and between the mixed runs in each session.

At each session, the experimental paradigm was performed in the MRI scanner after an initial T1-weighted scan during which participants watched a muted cartoon. The PC group followed the same protocol for their sessions A and D, while sessions B and C only included structural scans. For comparability, children who performed the task in the MRI simulator also watched a muted cartoon accompanied by scanner noise before they completed the task. Performance in this task-switching paradigm did not differ between children in the MRI and MRI simulator group at any of the sessions. Furthermore, the paradigm elicited reliable mixing and switch costs in accuracy and RT across all groups and sessions (see Supplementary Tables 7–8).

### 2.4 Behavioral analyses

Trials with response times (RTs) below 200 ms and above 3000 ms (i.e., responses given more than 1000 ms after the end of the stimulus presentation), and trials with no responses, were excluded from analyses. Based on these trimming criteria, of the 294 trials for each session (not including the first trial of each of the three runs), we excluded very few trials (on average the following number of trials per subject per session: M_A_ = 4.34 (SD_A_ = 4.83), M_B_= 3.34 (SD_B_ = 4.03), M_C_ = 3.44 (SD_C_ = 3.75), M_D_ = 3.48 (SD_D_ = 3.93)).

Deviating from the preregistration, we conducted the linear mixed models on trial-level data as opposed to aggregated participant-level data, allowing us to account for variance in performance within participants (Barr et al. 2013). All models included a random intercept of participant and random slopes of condition, task rule (face vs. scene vs. object), and session. All analyses were performed using Bayesian linear mixed models with the *brms* package in R (Bürkner 2017). Reported effects are based on 95% credible intervals (CI), meaning that we can make a statement with 95% probability (cf. Bürkner 2017; see also Morey et al. 2016). We started with a full model that included all interactions between fixed effects and compared this to models with fewer interaction terms, using leave-one-out cross-validation in the *loo* package (Vehtari et al. 2022). Across all analyses, the model including all interaction terms outperformed the models with fewer interaction terms (see Supplementary Table 3). There were no differences in performance among the three groups at session A prior to training (all 95%-CI included zero).

#### 2.4.1 Trajectories of change across all four sessions

To test for differences in the trajectories of change between the two training groups in the course of training, we compared accuracy and RTs between the SW and SI groups across all four sessions. Change across more than two sessions is not necessarily linear; to describe these trajectories as precisely as possible, we thus opted to include both a linear and a quadratic effect of session. Group (SW vs. SI), session (0–3), and condition (single vs. repeat vs. switch) were modeled as fixed effects. Additionally, we included the following three-way interactions: (1) group, condition, and the linear effect of session and (2) group, condition, and the quadratic effect of session. The models also included a random intercept of participant and random slopes of condition, task rule (face vs. scene vs. object), and session.

#### 2.4.2 Training-related changes from pre- to post-training

To evaluate whether changes were indeed related to training, we examined changes in accuracy and RTs in the SW and SI groups relative to the PC group from the pre-test (session A) to post-training (sessions D). Deviating from the preregistration, we included the SW and SI groups separately in these models, as opposed to combining them, to capture differences for each training group. More specifically, group (SW, SI, and PC, with PC as reference level), session (A vs. D), and condition (single vs. repeat vs. switch) were modeled as fixed effects, allowing for interactions among them.

### 2.5 fMRI data acquisition and preprocessing

As reported by Schwarze et al. (2023), structural and functional MR images were collected on a 3-Tesla Siemens Tim Trio MRI scanner. Functional runs consisted of 230 whole-brain echo-planar images of 36 interleaved slices (TR = 2000 ms; TE = 30 ms; 3 mm isotropic voxels; 32-channel head coil). The imaging protocol included functional imaging during the performance of the task-switching paradigm on all four sessions for the SW and SI group and on the first and last MRI session for the PC group. Structural MRI scans were acquired for all groups at all four sessions (220 slices; 1 mm isotropic voxels; TR = 4500 ms; TE = 2.35 ms; FoV = 160 ×198 x 220).

Preprocessing was performed using fMRIprep (Version 20.2.0; Esteban et al. 2019). For a detailed description, see the fMRIprep documentation (https://fmriprep.org/en/stable/). Briefly, functional images were co-registered to individual anatomical templates using FreeSurfer (Greve and Fischl 2009). To define the ROIs at session A, the anatomical template was created from anatomical scans of session A to limit the impact of images acquired after the start of training. For the data used in the GLMs for parameter extraction, the anatomical template was created from anatomical scans across all sessions, removing scans that were of poor quality based on the MRIQC classifier (Version 0.15.2; Esteban et al. 2017) and additional visual inspection. Images were slice-time corrected (using AFNI; Cox and Hyde 1997), realigned (using FSL 5.0.9; Jenkinson et al. 2002), resampled into MNI152NLin6Asym standard space with an isotropic voxel size of 2 mm, and finally spatially smoothed with an 8mm FWHM isotropic Gaussian kernel using SPM12 (Functional Imaging Laboratory, University College London [UCL], UK).

### 2.6 fMRI data analysis

#### 2.6.1 General linear models (GLM)

GLM analyses were performed using SPM12 software (Functional Imaging Laboratory, UCL, UK). For each participant, we estimated an event-related GLM for each session. Each stimulus presentation was coded as an event with zero duration, and convolved with a canonical hemodynamic response function. Separate regressors were included for correct single, correct repeat, and correct switch trials. Incorrect trials, trials with no responses, trials with extreme RTs (below 200 ms or above 3 s), and the first trial of each run were modeled in a nuisance regressor. Data were high-pass filtered at 128 s. To minimize head motion artifacts, we included the amount of framewise displacement per volume in mm (Power et al. 2012), realignment parameters (three translation and three rotation parameters), and the first six anatomical CompCor components (as provided by fMRIprep; Behzadi et al. 2007) as regressors of no interest. The first five volumes of each run were discarded to allow for stabilization of the magnetic field. Temporal autocorrelations were estimated using first-order autoregression.

To identify regions that showed activation associated with mixing demand, we compared activation on repeat trials in mixed blocks to single-block trials (repeat > single) across all three groups at session A (N = 89; voxel-level p < .001, FDR-corrected p < .05 at the cluster level). To identify regions modulated by switch demand, we compared activation on switch to repeat trials in the mixed blocks (switch > repeat) across all three groups at session A (N = 89; voxel-level p < .001, FDR-corrected p < .05 at the cluster level).

#### 2.6.2 ROI definition and analyses

ROIs were defined on the repeat > single contrast and switch > repeat contrast across all children at session A (described above). To ensure anatomical specificity, we anatomically restricted activation-based ROIs in the dlPFC, SPL, and precuneus using the middle frontal gyrus, SPL, and precuneus regions of the Harvard-Oxford atlas, respectively (thresholded at 30%; Makris et al. 2006). The IFJ was restricted based on coordinates from a meta-analysis of task switching (Derrfuss et al. 2005), as no anatomical mask is available for this functionally defined region associated with task switching (cf. Richter and Yeung 2014). We extracted activation parameters for these ROIs using Marsbar (Brett et al. 2002).

We performed Bayesian linear mixed models to investigate whether activation in the ROIs changed with training and differed between training groups. First, we explored trajectories of training-related change in activation across all four sessions in the two training groups. Models included fixed effects of group (SW vs. SI), session – modeled as a factor with four levels (A, B, C, D), and condition. Condition included repeat vs. single for models of activation in ROIs defined by the repeat > single contrast, and switch vs. repeat for activation in ROIs defined by the switch > repeat contrast. Random intercepts of participant and random slopes of session were also included in all models.

In a second step, to test whether changes in task-related activation were training-related, we compared the SW and SI groups to the PC group on sessions A and D, for which data from all groups was available. Models included fixed effects of group (SW vs. PC, SI vs. PC), session (A vs. D), and condition (repeat vs. single; switch vs. repeat) and their interactions, as well as random intercepts of individuals and random slopes of session. Results of these analyses are included in the main results for ROIs that showed changes in the two training groups across all four sessions; detailed model outputs for all ROIs are reported in Supplementary Tables 9 and 11.

To test for training-related changes in the adult task-switching network, we additionally defined ROIs from 53 adults (20–30 years) who performed the same task-switching paradigm at a single timepoint and did not undergo any training (see Schwarze et al. 2023 for details and comparison of children’s pre-training performance, activation, and connectivity to this group of adults). Results for these preregistered ROI analyses are reported in Supplementary Tables 4–6, and were generally consistent with the results reported below.

#### 2.6.3 Whole-brain longitudinal analyses

Since the ROI-based analysis is biased towards the activation patterns observed in session A, we additionally performed longitudinal whole-brain analyses to test for training-related changes outside the ROIs, as well as how these differed between groups. We constructed mixed ANOVAS in SPM with group as a between-participant factor and session as a within-participant factor. Specifically, we tested for differences between the SW and SI groups and the PC group comparing sessions A and D, and for differences between the SW and SI groups across all four sessions. The input contrast images included the repeat > single contrast to investigate changes in modulation of activation with mixing demand, and the switch > repeat contrast for changes in modulation of activation with switch demand. There were no significant clusters showing changes in activation with training at the predefined threshold (p < .001, uncorrected), neither when comparing sessions A and D nor when testing for any effects across all sessions.

#### 2.6.4 Psychophysiological interactions

To examine training effects in task-related functional connectivity, we conducted gPPI (generalized psychophysiological interaction) analyses (McLaren et al. 2012) using the CONN toolbox (Version 20b; Whitfield-Gabrieli and Nieto-Castanon 2012). gPPI can be used to model how connectivity strength differs between conditions, thus making it possible to investigate how brain networks are flexibly adapting to task demands. gPPI parameters were estimated separately for each condition, that is, correct single, correct repeat, and correct switch trials (McLaren et al. 2012). The main effect of the three conditions and the nuisance regressors from the activation GLM were regressed out of the fMRI timeseries before analysis. We calculated ROI-to-ROI gPPI for connections among ROIs associated with mixing demand, identified by the repeat > single contrast (i.e., bilateral IFJ, bilateral SPL, and left dlPFC). In a separate but identical model, we calculated ROI-to-ROI gPPI for connections among the ROIs associated with switch demands, identified by the switch > repeat contrast (i.e., left IFJ, bilateral SPL, bilateral precuneus).

The gPPI models provided two connectivity parameters for each connection between any two ROIs representing connectivity estimates in both directions. Therefore, we first tested whether the direction had an effect on the connectivity parameter. We modeled estimated connectivity for each connection with a linear mixed model including the direction, condition, session, and training group as fixed effects, allowing for all interactions and including random intercepts of individuals and random slopes of session. We compared this model to one without any interactions involving seed region. As model comparisons indicated better fit for models without interaction effects of seed region, we averaged parameters across directions for each connection to be used in the subsequent analysis.

Individuals’ condition-specific gPPI parameters (averaged across directions) for each connection and session were analyzed using Bayesian linear mixed models to examine differences in the changes of these parameters between the SW and SI groups. Specifically, mirroring the models of activation above, we tested whether connectivity values among the ROIs associated with mixing demands (defined on the basis of the repeat > single contrast) changed across sessions (i.e., session B, C, and D compared to A as the reference level). In addition to the fixed effect of session, models included fixed effects for group (SW vs. SI) and condition (repeat vs. single) and random intercepts of participant and connection, as well as random slopes for session. For the connectivity parameters among the ROIs associated with switch demand, we tested the same model, except that the condition levels consisted of switch vs. repeat. Matching the activation analyses, we subsequently tested whether changes in task-related connectivity were training-related by comparing the SW and SI groups to the PC group on sessions A and D.

Additionally, to characterize whether key task-switching regions changed connectivity to brain regions outside of the ROIs defined by brain activation, we analyzed seed-to-voxel PPIs. Here, we used a seed in the left IFJ and in the left SPL, based on their prominent roles in task switching (e.g., Kim et al. 2012; Richter and Yeung 2014) and our analyses of age differences in task switching between children and adults (Schwarze et al. 2023). Results of these seed-to-voxel analyses and further preregistered analyses, including the association among training-related changes in performance, activation, and connectivity are reported in Supplementary Results 1–3. We did not observe any associations between performance and neural measures in any of these analyses.

## 3. Results

### 3.1 Training-related changes in task-switching performance

#### 3.1.1 Overview of analytic approach

We expected that children in the SW group, who engaged in more task switching during training, would show greater and/or earlier improvements in task-switching performance than children in the SI group, who performed less task switching during training. To test this hypothesis, we compared the trajectories of changes across all four laboratory sessions in the SI and SW group by predicting accuracy and RTs by the fixed effects of group (SW vs. SI), condition (single, repeat, switch), the linear and quadratic effects of session, and their corresponding interactions. The inclusion of the quadratic effect of session allowed us to investigate more nuanced training-related changes beyond monotonic increases, for example whether improvements in performance slowed down or sped up towards the end of training.

While the models included all three conditions simultaneously, below we first describe the results including the condition comparisons pertaining to mixing costs (repeat vs. single) followed by the condition comparisons pertaining to switch costs (switch vs. repeat), considering how the costs in accuracy and RTs changed differentially between the training groups. For each of these sections, we performed additional control analyses to check whether changes were indeed related to training. Here, we compared change in the two training groups between sessions A and D to change in the PC group, which did not undergo any training. To simplify the description of results, we only report effects of session and interactions with session; complete model outputs can be found in Supplementary Tables 7 and 8.

#### 3.1.2 Training-related improvements in accuracy

Overall accuracy increased with training (linear effect of session: est. = 0.62, 95%-CI: 0.46, 0.78; quadratic effect of session: est. = –0.22, 95%-CI: –0.26, –0.18; see Supplementary Tables 7 and 8 for complete results). These overall changes were qualified by interactions with condition and group indicating changes in mixing and switch costs, which further differed between training groups. We therefore consider these changes in mixing and switch costs separately next.

##### Mixing costs

Accuracy mixing costs decreased rapidly at the beginning of training in both training groups, with slower decreases towards the end of training (Figure 2A): linear and quadratic changes in accuracy were more pronounced for repeat than single trials (linear effect of session x condition [single vs. repeat]: est. = 0.50, 95%-CI: 0.29, 0.72; quadratic effect of session x condition [single vs. repeat]: est. = –0.13, 95%-CI: –0.20, – 0.06). Of note, children in the SW group maintained high accuracy on repeat trials towards the end of training, thereby maintaining lower mixing costs compared to children in the SI group (linear effect of session x condition [single vs. repeat] x group [SW vs. SI]: est. = – 0.48, 95%-CI: –0.80, –0.16; quadratic effect of session x condition [single vs. repeat] x group [SW vs. SI]: est. = 0.11, 95%-CI: 0.00, 0.22).

**Figure 2:**
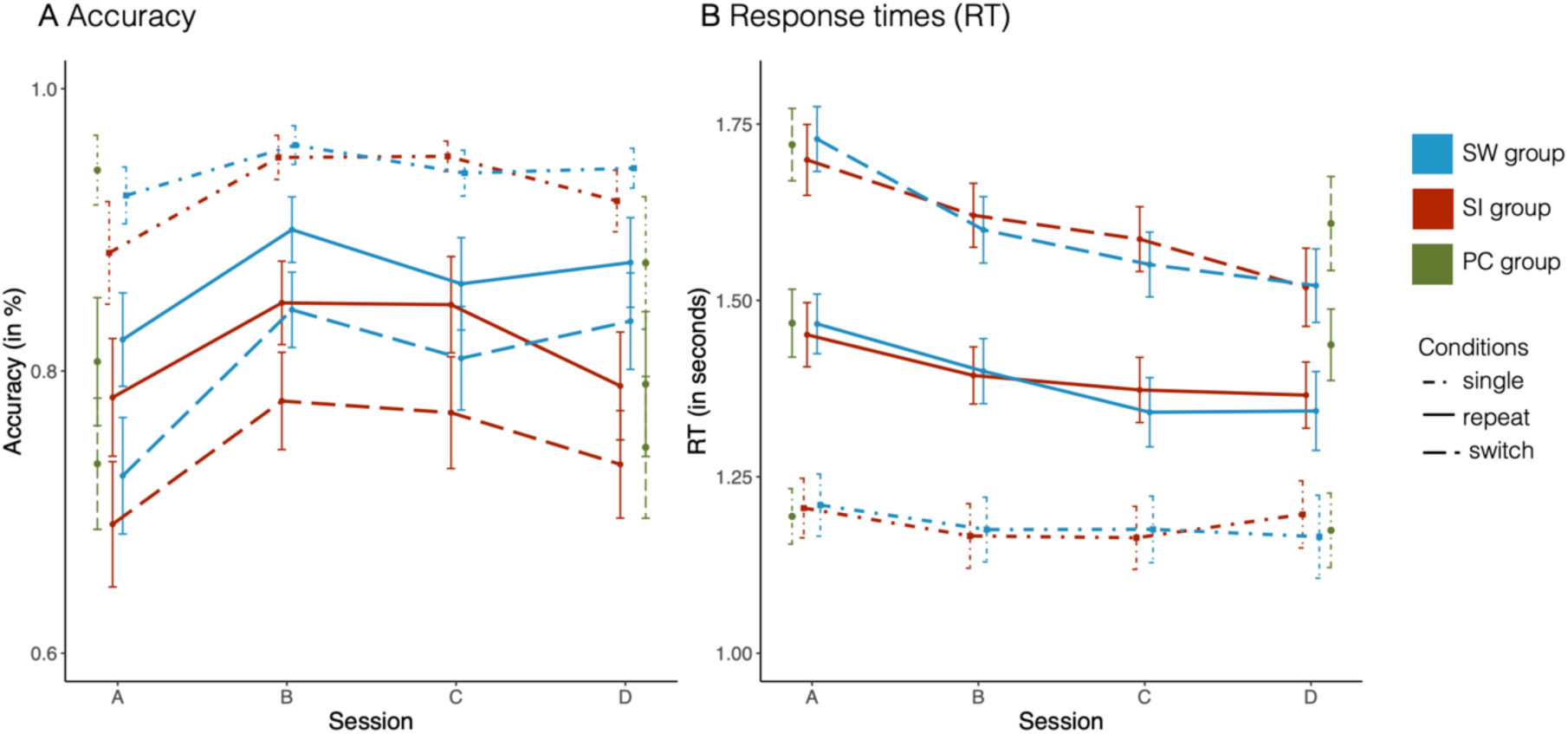
Training-related changes in performance. Accuracy (A) and response times (B) for each of the three conditions: single trials (dot-dashed line), repeat trials (solid line), and switch trials (dashed line). The SI group is shown in red, the SI group in blue, and the PC group in green. Note that the figure shows aggregated data while models were conducted at trial level. Error bars denote 95%-confidence intervals.

Comparisons to the PC group confirmed the selective mixing-cost improvements in the SW group: Compared to the PC group, accuracy mixing costs decreased in the SW group due to greater accuracy improvements on repeat trials than on single trials from pre- to post-test (session [D vs. A] x condition [single vs. repeat] x group [SW vs. PC]: est. = 0.48, 95%-CI: 0.12, 0.84). Even though mixing costs did not change from session A to session D in the SI group, at session D the SI group showed larger mixing costs than the PC group due to declines in single-trial accuracy in the PC group (session [D vs. A] x condition [single vs. repeat] x group [SI vs. PC]: est. = 0.98, 95%-CI: 0.66, 1.30).

##### Switch costs

There were no changes in accuracy switch costs (all 95%-CI include zero) across any of the groups in the present study.

#### 3.1.3 Training-related improvements in RT

Overall RTs decreased with training (linear effect of session: est. = –0.07, 95%-CI: –0.09, – 0.05; quadratic effect of session: est. = 0.02, 95%-CI: 0.01, 0.02). These overall changes were qualified by interactions with condition and group, indicating changes in mixing and switch costs, which further differed between the training groups. We therefore turn to these changes in mixing and switch costs separately next.

##### Mixing costs

RT mixing costs decreased in both training groups (Figure 2B), but showed distinct trajectories of condition-specific change between the SI and SW groups (linear effect of session x condition [single vs. repeat] x group [SW vs. SI]: est. = –0.05, 95%- CI: –0.08, –0.02; quadratic effect of session x condition [single vs. repeat] x group [SW vs. SI]: est. = 0.02, 95%-CI: 0.01, 0.03). In the SW group, RT mixing costs decreased because children became progressively faster on repeat trials, indicating performance improvements. In contrast, in the SI group, responses on single trials became faster initially but slowed down again towards the end of training, returning to baseline levels. The comparison to the PC groups showed that reductions in RT mixing costs were more prominent in the SI and SW groups than the PC group (session [D vs. A] x condition [single vs. repeat] x group [SI vs. PC]: est. = 0.06; 95%-CI: 0.02, 0.09; session [D vs. A] x condition (single vs. repeat) x group [SW vs. PC]: est. = 0.04; 95%-CI: 0.00, 0.08).

##### Switch costs

While RTs on repeat trials decreased earlier during training, RTs on switch trials declined only towards the end of training – but to a greater extent – resulting in RT switch cost decreases towards the end of training in both training groups (linear effect of session x condition [switch vs. repeat]: est. = 0.03, 95%-CI: 0.01, 0.05; quadratic effect of session x condition [switch vs. repeat]: est. = –0.02, 95%-CI: –0.02, –0.01). This pattern was more prominent in the SI group than the SW group (linear effect of session x condition [switch vs. repeat] x group [SW vs. SI]: est. = –0.05, 95%-CI: –0.08, –0.01; quadratic effect of session [switch vs. repeat] x condition x group [SW vs. SI]: est. = 0.02, 95%-CI: 0.01, 0.03). However, the decreases in RT switch costs between pre- and post-test in the training groups were comparable to the PC group as evident in the interaction of session and condition across all three groups (session [D vs. A] x condition [switch vs. repeat]: est. = – 0.06; 95%-CI: –0.09, –0.03), and the lack of group differences therein (all 95%-CI included zero), suggesting that the decrease in RT switch costs was not related to training.

#### 3.1.4 Summary of behavioral results

In both training groups, accuracy and RT mixing costs improved quickly in the first training sessions, followed by slower improvements towards the end of training. Group differences associated with varying amounts of task-switching practice became particularly prominent in the last training sessions, when the SW group maintained the initial training benefits, while performance in the SI group returned to similar performance levels as pre-training. Thus, contrary to our expectations, the two training groups showed a similar extent and speed of improvements but differed in how their improvements were maintained over the later phase of training. Comparisons to a passive control group at pre- and post-test offer further support that these changes in mixing costs were indeed related to intensive task-switching practice. In contrast, we found no evidence for training-related reductions in accuracy or RT switch costs.

### 3.2 Analyses of task-related activation and connectivity

To investigate training-related changes in the neural processes associated with mixing costs and switch costs, we examined task-related activation in and connectivity among the ROIs associated with greater activation on repeat than single and switch than repeat trials, respectively. These analyses aimed to distinguish between two hypotheses: First, we hypothesized that frontoparietal activation decreases with training, indicating more efficient processing in these regions. Alternatively, we hypothesized that children’s activation becomes more adult-like with training, such that frontoparietal activation increases more steeply with mixing demand (i.e., on repeat relative to single trials) and with switch demand (i.e., on switch relative to repeat trials). With regard to task-based connectivity, we expected that (1) connectivity among frontoparietal regions would increase with training and (2) differences between conditions would become smaller as connections become more efficient with training.

Our analyses proceeded in two steps. First, we tested whether activation and connectivity changed at any of the sessions B, C, D relative to the pre-test session A for the two training groups (i.e., SW vs. SI). Second, to test whether these changes were indeed related to training, we compared changes in the training groups to the PC group, focusing on sessions A and D where data from all groups were available. To simplify the description of results, we report effects involving session and interactions of session with group (see Supplementary Tables 9–12 for an overview of all effects).

#### 3.2.1 Repeat vs. single: Training-related changes in activation and connectivity

Whole-brain activation across all children at session A revealed stronger activation for repeat > single trials in multiple frontal and parietal regions, including the bilateral IFJ, bilateral SPL, and left dlPFC (Figure 3A). Accordingly, we investigated training-related change in activation and ROI-to-ROI connectivity on repeat and single trials in the following ROIs: left and right IFJ, left and right SPL, and left dlPFC.

**Figure 3:**
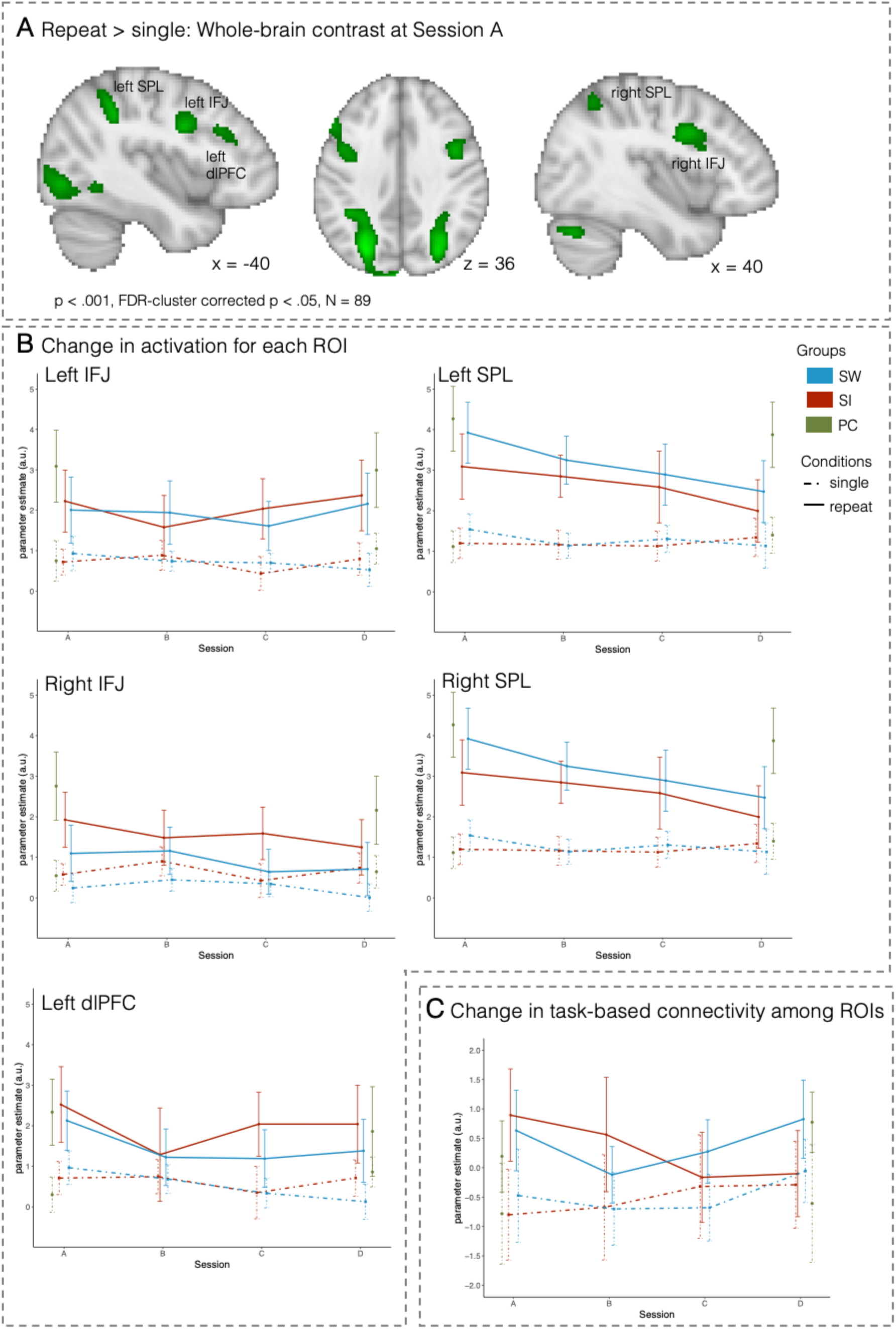
Training-related changes in activation and connectivity associated with mixing costs. (A) Brain regions showing greater activation on repeat than on single trials at session A across all children (N = 89; p < .001, FDR-cluster corrected p < .05). (B) Change in activation for each ROI. (C) Change in connectivity (i.e., PPI parameters across all connections) among these ROIs. The SW group is shown in blue, the SI group is shown in red, and the PC group is shown in green (for sessions A and D only). Error bars denote 95%-confidence intervals.

##### 3.2.1.1 Reduced frontoparietal activation with training in the SW group

In the right SPL (Figure 3B), activation on repeat trials decreased in both groups over time, resulting in a diminishing difference between single and repeat trials that became evident at session D (est. = –1.05; 95%-CI: –2.09, –0.03). While a descriptively similar pattern was evident in the left SPL, the data did not support the effect with 95% probability. Comparison to the PC group for sessions A and D challenges the interpretation of these changes as training-related: neither the SW nor the SI group differed reliably from the PC group at session D (all 95%-CI include zero).

Relative to single trials, repeat-trial activation in the left dlPFC (Figure 3B) decreased across both training groups from session A to B (session [B vs. A] x condition [repeat vs. single]: est. = –1.40; 95%-CI: –2.54, –0.23), consistent with the mixing costs improvement observed in performance. This effect was no longer evident at sessions C and D (95%-CIs included zero), when the difference between activation on repeat and single trials did not differ from session A in either group (i.e., 95%-CIs include zero).

Comparison to the PC group across sessions A and D indicated that the SW group showed more pronounced decreases in dlPFC activation across both repeat and single trials from session A to D (session [D vs. A] x group [SW vs. PC]: est. = –1.48; 95-%CI: –2.50, – 0.31). The SI group did not show any activation differences relative to the PC group (session [D vs. A] x group [SI vs. PC]: est. = –0.54; 95-%CI: –1.66, 0.53). A descriptively similar pattern, albeit not at 95% probability, became evident in the left IFJ.

Thus, dlPFC showed differential training-related decreases in activation across groups. The effects in dlPFC were observed in two phases. First, after three weeks of training, activation on the more demanding repeat trials decreased for both training groups. Second, at the end of training, the *difference* between repeat and single trials had returned to the pre-training level, but in the SW group, this was due to decreasing activation on single trials, like repeat trials, with training, resulting in this group’s overall decreased activation, while in the SI group, this was due to repeat-trial activation increasing again. Thus, the results in the SW group are consistent with previous studies in adults that have reported decreasing frontoparietal activation with training.

##### 3.2.1.2 Decreased task-based connectivity in the SI training group

To test whether task-based connectivity among the ROIs (i.e., left IFJ, left SPL, left dlPFC, right IFJ, right SPL) showed training-related changes, we analyzed connection-specific PPI parameters for the effects of session (B, C, and D compared to A) and interactions with group (i.e., differences between SW and SI groups).

Across sessions, connectivity was greater for repeat than for single trials (repeat vs. single: est. = 1.91; 95%-CI: 1.46, 2.38), indicating that regions interacted more closely in response to mixing demands. As shown in Figure 3C, this condition difference was influenced by training. Specifically, connectivity for the repeat condition decreased with training in both groups (condition [repeat vs. single] x session [B vs. A]: est. = –0.75; 95%- CI: –1.46, –0.05; C vs. A: est. = –2.17; 95%-CI: –2.85, –1.49; D vs. A: est. = –1.57; 95%-CI: –2.25, –0.88), with especially pronounced decreases in the SI group towards the end of training (condition [repeat vs. single] x group [SW vs. SI] x session [C vs. A]: est. = 2.14; 95%-CI: 1.23, 3.05; [D vs. A]: est. = 1.41; 95%-CI: 0.47, 2.36). Comparison to the PC group for sessions A and D indicated that this late change was specific to the SI group: compared to the PC group, the SI group showed lower connectivity on repeat trials at session D compared to A (condition [repeat vs. single] x group [SI vs. PC] x session [D vs. A]: est. = – 1.58; 95%-CI: –2.60, –0.58). The SW group did not differ from the PC group at session D (95%-CI includes zero).

In summary, contrary to the first part of our hypothesis, we did not observe stronger connectivity with training. In line with the second part of our hypothesis, connectivity among frontoparietal ROIs during repeat trials decreased with training across both groups, suggesting less upregulation of connectivity in the mixed- vs. single-task condition over time. Towards the end of training (i.e., sessions C and D) this decrease was particularly pronounced for the SI group relative to the SW group.

#### 3.2.2 Switch vs. repeat: Training-related changes in activation and connectivity

Whole-brain activation across all children at session A indicated increased activation for switch than for repeat trials in the left IFJ, bilateral SPL, and bilateral precuneus. Accordingly, we investigated training-related changes in activation and ROI-to-ROI connectivity on switch and repeat trials in the following ROIs: left IFJ, left and right SPL, and left and right precuneus (Figure 4A).

**Figure 4:**
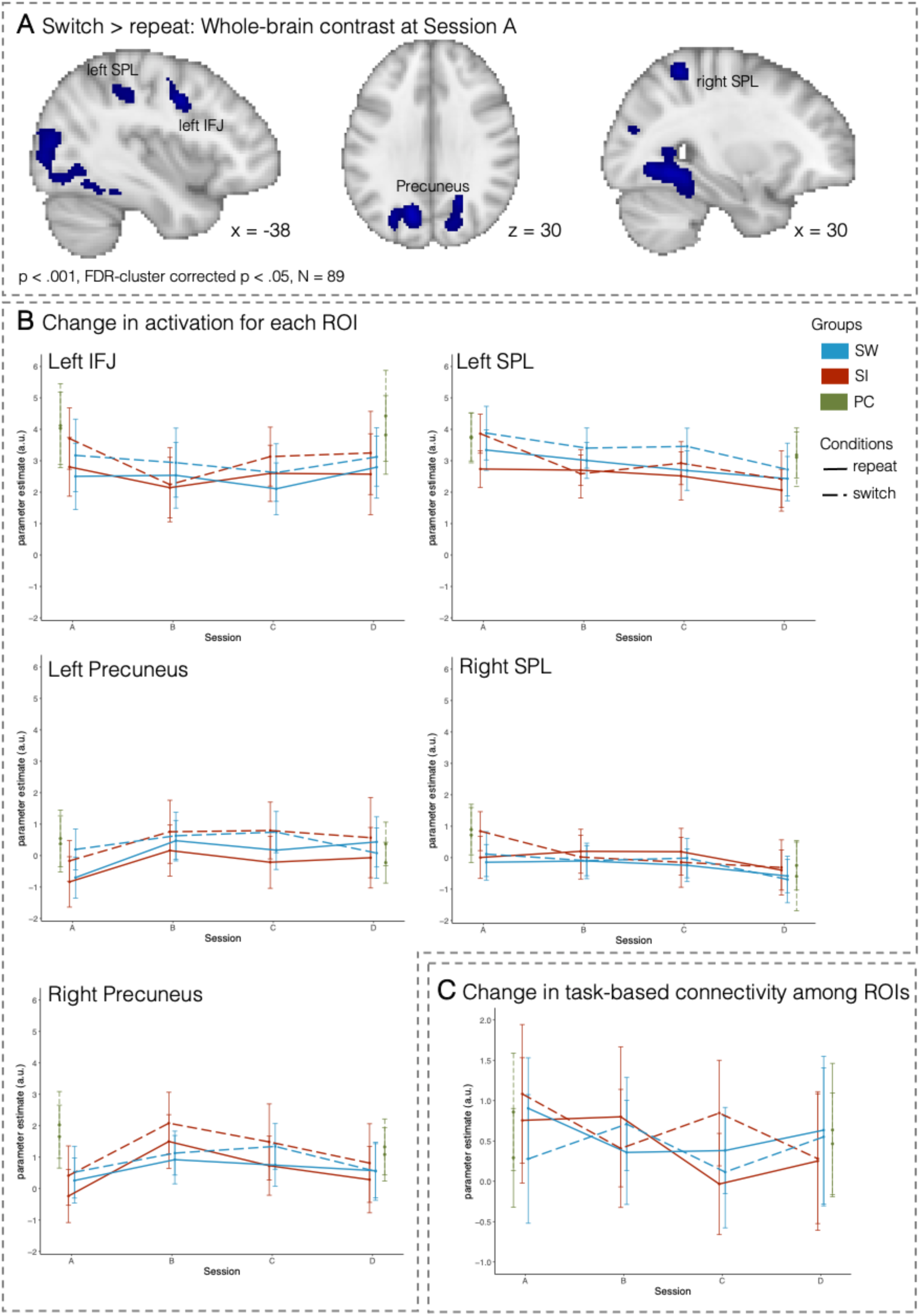
Training-related changes in activation and connectivity associated with switch costs. (A) Brain regions showing greater activation on switch than on repeat trials at session A across all children (N = 89; p < .001, FDR-cluster corrected p < .05). (B) Change in activation for each ROI. (C) Change in connectivity (i.e., PPI parameters across all connections) among these ROIs. The SW group is shown in blue, the SI group in red, and the PC group in green (for sessions A and D only). Error bars denote 95%-confidence intervals.

##### 3.2.2.1 Transient changes in switch-related brain activation with training

Across both training groups (Figure 4B), we observed decreased activation in multiple ROIs from session A to B across switch and repeat trials (left precuneus: est. = 0.99; 95%-CI: 0.17, 1.82; right precuneus: est. = 1.22; 95%-CI: 0.39, 2.04), and specifically for switch trials (left SPL: session [B vs. A] x condition [switch vs. repeat]: est. = –1.07; 95%-CI: –1.83, – 0.33). These changes returned to baseline towards the end of training (i.e., sessions C or D vs. A: all 95%-CI include zero). However, the interpretability of these effects as related to training is limited: while activation in the left SPL and right precuneus did not differ from the control group at session D (all 95%-CI include zero), results in the left precuneus were driven by overall group differences in activation between the training groups and the control group (Figure 4B; group [SI vs. PC]: est. = –1.26; 95%-CI: –2.32, –0.20; group [SW vs. PC]: est. = –1.05; 95%-CI: –2.07, –0.03).

The right SPL showed decreases in activation on switch trials over time (session [C vs. A] x condition [switch vs. repeat]: est. = –0.68; 95%-CI: –1.30, –0.09; session [D vs. A] x condition [switch vs. repeat]: est. = –0.66; 95%-CI: –1.31, –0.01). However, comparisons to the PC group did not show evidence that changes in the training groups differed from the PC group (all group x session interactions 95%-CI included zero).

To summarize, the ROIs associated with switching demands within mixing blocks predominantly showed transient changes in the beginning of training that returned to initial levels in the later training sessions. Overall, a lack of differences to the PC group at session D cautions against an interpretation of this decrease as related to training.

##### 3.2.2.2 Inconclusive results regarding changes in task-based connectivity with training

We next analyzed connection-specific PPI parameters among left IFJ, left SPL, left precuneus, right SPL, and right precuneus for the effects of session (B, C, D compared to A) and interactions with group (SW vs. SI group).

As shown in Figure 4C, connectivity for switch trials decreased from session A to B in the SI group, while the SW group showed an increase of connectivity for switch trials and a decrease for repeat trials (session [B vs. A] x group [SW vs. SI] x condition [switch vs. repeat]: est. = 1.77; 95%-CI: 0.76, 2.77). However, we did not observe clear differences in connectivity between switch and repeat trials across the two groups independent of training. Thus, the observed group differences cannot be meaningfully interpreted as training-related changes in adaptively adjusted connectivity based on switch demands.

## 4. Discussion

In the present study we investigated the trajectories of behavioral and neural changes in 8– 11-year-olds as a function of different amounts of task-switching training. To this end, we analyzed task performance, fMRI activation, and task-based functional connectivity before and after training, as well as twice during the nine-week training phase.

Comparing the pre- and post-training sessions, we observed training-related decreases in mixing costs, but not in switch costs. While RT mixing costs decreased in both training groups, accuracy mixing costs decreased only in the group with the higher amounts of task switching during training. These findings corroborate and extend previous findings demonstrating that task-switching performance in children, especially with regard to mixing costs, can be improved with intensive training (Karbach and Kray 2009; Kray, Karbach, Haenig, et al. 2012; Zinke et al. 2012; Dörrenbächer et al. 2014; Karbach et al. 2017).

By leveraging additional laboratory assessments during training, we provide a more detailed picture of how the different training schedules influenced the trajectories of performance change. Mixing costs improved rapidly after the start of training. Contrary to our hypotheses, the two training groups showed similar improvements in this early phase of training. Instead, group differences emerged in the later phase of training: the intensive task-switching group maintained the training benefits from the first phase, whereas the group with lower amounts of task-switching training showed performance declines towards the end of training, returning to baseline levels; possible reasons for this decline are discussed below.

At the neural level, we observed different patterns of activation changes associated with mixing demands (i.e., repeat vs. single trials) and switch demands (i.e., switch vs. repeat trials). Compared to the passive control group, activations for both single and repeat trials decreased in the left dlPFC in the intensive task-switching group, but not in the intensive single-tasking group. Activation decreases in the dlPFC first became evident on the more demanding repeat trials and were followed by decreased activation on single trials only later in the course of training, resulting in similar condition *differences* between pre- and post-training assessments. While the SPL showed changes in activation related to mixing and switch costs, a lack of differences to the control group cautions the interpretation of these findings as training-related and we thus refrain from further discussing it. In parallel, connectivity among frontal and parietal regions decreased with training for repeat (but not single) trials, with more pronounced changes in the intensive single-tasking group than in the intensive task-switching group. Brain regions specifically associated with switch demands showed rapid training-related activation changes early in training, but returned to baseline by the end of training.

Taken together, more intensive task-switching training led to rapid improvements in mixing costs that were maintained over the course of training. The accompanying neural analyses indicate that intensive task-switching training was associated with decreases in task-related activation in the dlPFC as well as in connectivity among prefrontal and parietal regions, presumably indicating more efficient task processing with training.

### 4.1 Training improves efficiency of processing in frontoparietal regions

Both training groups showed decreased RTs, especially on mixed blocks, suggesting faster processing of task demands when multiple task rules had to be maintained, monitored, and reconfigured. Faster or more efficient rule processing with multitasking training has been proposed by Dux and colleagues (2009), based on decreased activation in lateral PFC, a region commonly associated with rule processing during task switching (Richter and Yeung 2014). Interestingly, our data of activation changes in the dlPFC suggests that rule processing in the more demanding condition (i.e., repeat compared to single trials) improved first, followed by more efficient processing of rules during single tasking.

Previous studies in adults have associated training-related activation decreases with increased efficiency of rule processing (Kelly and Garavan 2005; von Bastian et al. 2022). Poldrack (2000) suggested that training-related activation decreases result from more specific recruitment of neural circuits, more precise neural representations of task sets that enable more efficient processing of these task sets, or both. Accordingly, Garner and Dux (2015) have provided evidence for the role of representations based on multivariate activation patterns: with dual-task training, they observed that –along with decreases in univariate activation– task-set representations became more distinct, and that increased neural distinctiveness was associated with improved performance. The authors concluded that the increased precision of task-set representations aids their processing in prefrontal brain regions during dual-tasking and thus enables increased efficiency evident in reduced activation of prefrontal regions (see also Dux et al. 2009). Such improved distinctiveness in representations would be especially beneficial within mixed contexts (i.e., on repeat compared to single trials), where multiple task-set representations need to be continuously maintained. Future studies using multivariate methods to investigate whether task-switching training results in more distinct neural representations of task sets in children can help to better understand the activation decreases observed here (Garner and Dux 2015). The authors further showed that activation peaked earlier after training, suggesting that faster recruitment of neural circuits contributed to the training-related performance improvements as well (Dux et al. 2009). Thus, it is likely that both changes in representation and the speed with which a neural circuit is activated may contribute to activation decreases after training.

The present results extend the observation that increased efficiency underlies training-related improvements in executive functions from adulthood (cf. von Bastian et al. 2022) to late childhood, by demonstrating decreases in frontal activation that were accompanied by decreased mixing costs with intensive task-switching training. This observation is relevant in light of suggestions that training in children may speed up maturation with children becoming more adult-like in activation and connectivity patterns (Jolles et al. 2012; Jolles and Crone 2012). Our previous work (Schwarze et al. 2023) showed that the present task-switching paradigm elicited smaller upregulation of frontoparietal activation with increased task demands in (untrained) children compared to adults. This pattern indicated a potential for children’s activation patterns to become more adult-like by condition-specific increases in activation. For instance, we would have expected increased activation on repeat trials with training to match the pattern of condition differences in activation observed in adults performing the same paradigm. However, we did not observe such changes in any of the ROIs (see also Supplementary Results 3). Rather than showing more adult-like activation, children revealed a similar training-related change of decreasing PFC activation as previously reported in (trained) adults. Future studies are needed to elucidate how these training-related changes in neural processes depend on the targeted executive function (e.g., task-switching as opposed to working-memory training; Jolles et al. 2012; Astle et al. 2015) or the investigated age range (Rueda et al. 2005; Lee et al. 2022).

While the intensive single-task group did not show training-related changes in task-related activation at the end of training, they showed more pronounced changes in task-based connectivity. Specifically, connectivity among frontoparietal regions in the intensive single-task group decreased for repeat trials, resulting in smaller differences in connectivity on repeat compared to single trials at the end of training. These connectivity decreases seem counterintuitive in light of previous training studies showing connectivity increases in children at rest (Astle et al. 2015; Lee et al. 2022) and in adults during task performance (Kundu et al. 2013; Thompson et al. 2016). Contrary to these findings, connectivity decreases on repeat trials as observed here could indicate that frontoparietal connections were less strongly recruited on these relatively more demanding trials after training.

Note that it is unclear whether connectivity decreases in our study were beneficial or detrimental to task-switching performance, given that the performance of children in the intensive single-task training group returned to pre-training levels at the end of training. The observed decrease in task-based connectivity could reflect more localized processing that requires less recruitment of multiple regions in concert. Alternatively, baseline connectivity (i.e., independent of specific task demands) could have increased with training (cf. Astle et al., 2015; Lee et al., 2022), which would require smaller additional increases with task demands to reach a similarly strong connection as before training. Future studies combining resting-state and task-based connectivity measures would be able to disentangle these hypotheses.

### 4.2 Differences between training-related changes in mixing and switch costs and their associated activation

Training-related performance improvements in the present study were limited to decreases in mixing costs. The two training groups showed lower switch costs in response times only at the end of training. However, as the magnitude of change did not differ between the training groups and the passive control group, the decreases in switch costs most likely reflect test-retest effects rather than intensive training (cf. Bartels et al. 2010; Brahms et al. 2021). Moreover, most activation changes associated with switch costs (i.e., switch vs. repeat trials) were similar across training groups and relatively short lived: changes became evident at the second session but were no longer present at the end of training. As a result, switch-related activation in the training groups did not differ from the passive control group at the end of training, limiting the interpretation of the activation changes as training-related.

The lack of training-related decreases in switch costs (i.e., differences between switch and repeat trials) should be interpreted in light of the fact that RTs decreased on both switch and repeat trials at the beginning of training, followed by more substantial RT decreases specific to switch trials only towards the end of training. Decreased RTs on both switch and repeat trials could reflect a general improvement in the maintenance and management of multiple rules. Specifically, given that our paradigm included three different tasks, each with three possible S-R mappings as compared to mostly paradigms with two rules with two possible S-R mappings used in previous studies (e.g., Karbach and Kray 2009; Kray, Karbach, Haenig, et al. 2012; Zinke et al. 2012; Zuber et al. 2023), it may have provided substantial room for improvement across both switch and repeat trials resulting in training effects on mixing costs but not switch costs. This increased demand is consistent with the fact that children in the present study showed smaller switch costs than mixing costs prior to training (see Schwarze et al. 2023).

The different patterns of change observed for activation associated with mixing and switch costs can be considered within the framework of supply-demand mismatch of training-related neural plasticity (cf. Lövdén et al. 2010; Lövdén et al. 2020). According to this framework, training-related changes are expected to be particularly pronounced when the mismatch between current ability (i.e., supply) and task demands is large enough but not beyond reach. Since we previously reported larger age differences between children and adults for mixing costs than for switch costs before training (Schwarze et al. 2023), we can assume a greater mismatch between current ability and task demands for mixing costs than for switch costs in the present study. Due to the smaller mismatch for switch costs, the changes in switch-related activation observed early during training may not have been long lasting. Given that the gap in switching performance to adult levels is more pronounced at earlier ages, the supply-demand mismatch account would predict more substantial and longer-lasting changes in switch-related activation for younger children, who perform task switching with lower proficiency (e.g., in the 4–7 age range). Importantly, Lövdén and colleagues (Lövdén et al. 2020; Lövdén & Lindenberger 2019) posited that long-lasting training-related changes require changes in brain structure. Therefore, future studies are needed to investigate whether and how cognitive training is associated with changes in gray-matter structure and structural connectivity in children.

### 4.3 Return to baseline performance and activation in the SI group

While we had predicted slower or less extensive behavioral improvements and neural changes with lower dosages of task switching, the intensive single-tasking group showed behavioral and neural changes as quickly and almost as extensively as the intensive task-switching group. However, the intensive single-tasking group returned to baseline levels of performance and dlPFC activation by the end of training. There are different potential explanations for this return to baseline.

A key difference between the two training groups may be related to the need to track changes in task demands indicated by changes in context. The ability to track changes in contexts continues to develop in late childhood (Waskom et al. 2014; Frick and Chevalier 2023) and contributes to the development of self-directed control (Frick et al. 2022). In the present training paradigm, the demands for context tracking were greater for the intensive task-switching group, in which participants performed more mixed blocks during training and thus faced more frequent switches of the cue. During the training games, the cue was always presented along with the target stimulus, effectively rendering it the context of the stimulus. Thus, the intensive task-switching group not only learned the mappings between each stimulus and the corresponding response, but also more intensively practiced tracking the context in which these were presented, which was crucial for successful task performance. In turn, the improved ability to track the context might have enabled more efficient rule implementation in the dlPFC (Hyafil et al. 2009; Ruge et al. 2013) and thus reduced activation in this region to a greater extent in the intensive task-switching group than in the intensive single-tasking group.

Additionally, children in the two training groups may have learned the rule structure differently, based on their experience with the tasks during training. Mixed blocks require the application of the rules in a hierarchical manner: mappings between a stimulus and a response are nested within a cue indicating which stimulus is relevant. While the ability to identify and apply such hierarchical rule structures has been demonstrated in infants and toddlers (Werchan et al. 2015; Werchan et al. 2016), it is continually refined throughout childhood and adolescence (Kray, Karbach, and Blaye 2012; Unger et al. 2016). During the instruction phase of each training game, the hierarchical structure of each task was made explicit to both groups. Nonetheless, children in the intensive single-tasking group may have represented the mappings between stimulus and response separately from the cue, as the cue was only relevant at the beginning of a single-task block and then remained the same for the duration of this block. In other words, the intensive single-task training group may have represented all rules encountered during training separately. Towards the end of training, such flat rule representations could have resulted in increasing rule competition and higher performance costs. In contrast, the intensive task-switching training group needed to utilize the associations between cues, stimuli, and response to a greater extent during training, likely strengthening the associations amongst them, resulting in lower costs.

Finally, it is important to point out that the MRI paradigm was one of the training games and was performed on 5 out of 27 training sessions by all children, with the SI group performing mostly single-task blocks in this paradigm. Independent of tasks being mostly performed intermixed or separately, children in both groups therefore acquired extensive practice of these specific stimulus-response mappings, which might have contributed to the changes in performance, activation, and connectivity for this specific paradigm. Rule-specific learning might have also contributed to the differences between the two training groups: learning in the intensive task-switching group included longer intervals between retrieval of individual rules due to the mostly intermixed presentation of the rules; in parallel, the intensive single-tasking group underwent massed learning of the rules due to higher exposure to single-task blocks.

### 4.4 Limitations and future directions

We would like to acknowledge several important limitations of the present study. First, the sample size for the analyses of activation and connectivity is relatively small, limiting our ability to find smaller effects, especially for whole-brain analyses over time as they require complete datasets with currently available data-analysis pipelines. Thus, the reported training-related changes in activation and connectivity patterns should be interpreted with caution and seen as a starting point for further research. Additionally, changes in performance and brain function may happen at different time scales (Baykara et al. 2021). For example, the limited changes in activation associated with switch costs may be due to session B being too far into training to capture the potentially early onset of such changes. Different time scales of changes in behavior and brain function and the relatively small sample size might further contribute to the lack of association between individual differences in behavioral and neural change.

While we contrasted two main hypotheses regarding training-related change in task-related activation in children, we would like to acknowledge that the changes associated with each hypothesis might take place in parallel. Specifically, with training, children may show activation decreases, reflecting greater efficiency, and they might show increasing activation specifically in the more demanding condition, reflecting a shift towards more adult-like activation. If these processes take place simultaneously, we might observe a pattern manifested in a lack of change. Alternatively, if these processes happen on different time scales, they may together produce a U-shaped trajectory similar to the trajectory observed for the intensive single-tasking group in the present study. Future studies using shorter intervals between neuroimaging assessments during training can help elucidate how these two processes interact and whether complex patterns such as U-shaped trajectories could be caused by their interaction. Moreover, changes in fMRI activation need to be interpreted with caution, especially when inferring increased efficiency based on decreased activation, as commonly done in training studies. Future studies need to investigate how decreasing activation with training is associated with changes in brain metabolism to support the claim of increased efficiency (cf. Poldrack, 2015).

While we demonstrated that different amounts of task-switching training (i.e., intensive task switching vs. intensive single tasking) were associated with differential changes in performance and activation patterns, it seems worthwhile to investigate the effects of other training regimes in future work. Finally, we would like to stress that our goal was to specifically explore the neural basis of short-term plasticity in the neural system that supports task switching rather than to assess transfer of task-switching training to other cognitive functions, for which there is mixed evidence to date (see Diamond and Ling 2019 for a review). Future neuroimaging studies are needed to understand the neural mechanisms of transfer of training effects.

### 4.5 Conclusion

In this study, we showed that intensive task-switching training led to decreased mixing costs and reduced activation and connectivity in frontal regions in children aged between 8 and 11 years. By comparing different amounts of task switching during training, we were able to uncover the dynamics of performance improvements with varying demands. Improvements with task-switching training occurred quickly and were present with both lower and higher amounts of task-switching practice. However, task-switching demands played a key role for maintaining these initial benefits, such that children with intensive task-switching training were the ones who sustained higher task-switching performance at the end of training.

Our findings provide initial evidence on the ways in which processes associated with task-switching change with training at both behavioral and neural levels of analysis in late childhood, suggesting that cognitive training does not simply make children more adult-like but rather alters performance via increased efficiency of task processing in the lateral PFC. While intensive task-switching training resulted in decreases of activation for both single and repeat trials (though at a different pace), intensive single-task training was associated with decreases in connectivity for repeat trials.

Together, our findings reveal that different amounts of task-switching training over an extended period are associated with distinct neural change profiles. Future research can build on these findings by investigating which training regimes are most effective in promoting efficient task switching at different ages, with the aim of better understanding the interplay between maturational and experiential factors and timescales in the ontogeny of cognitive control.

## Acknowledgments

During the work on their dissertations, S.A.S. and N.K. were predoctoral fellows of the International Max Planck Research School on the Life Course (LIFE; http://www.imprs-life.mpg.de) at the Max Planck Institute for Human Development, Berlin, Germany. The authors thank Theodoros Koustakas for support with the analyses and Julia Delius for editorial assistance and helpful comments. Communication regarding this article should be directed to S.A.S, or Y.F..

