## Supplementary Material for "Intensive task-switching training and single-task training differentially affect behavioral and neural manifestations of cognitive control in children"

**Schwarze et al.**

### Supplementary Materials

| Content | Page |
|---|---|
| Supplementary Table 1. Details of training games | 3 |
| Supplementary Table 2. Details of trial distributions of training games | 7 |
| Supplementary Table 3. Model comparisons | 8 |
| Supplementary Table 4. Changes in activation across all sessions (A–D) in adult ROIs (repeat > single) | 9 |
| Supplementary Table 5. Changes in activation across all sessions (A–D) in adult ROIs (switch > repeat) | 10 |
| Supplementary Table 6. Details on ROIs defined on the adult comparison sample | 11 |
| Supplementary Table 7. Model outputs of performance (accuracy and response times) across all sessions (A vs. D) | 12 |
| Supplementary Table 8. Model outputs of performance (accuracy and response times) across all sessions (A–D) | 13 |
| Supplementary Table 9. Changes in activation between session A and D in all three groups (SI vs. MC, SW vs. MC) for repeat and single trials in the ROIs defined on the repeat > single contrast at session A. | 14 |
| Supplementary Table 10. Changes in activation across all sessions (A, B, C, and D) in the two practice groups (SI vs. SW) for repeat and single trials in the ROIs defined on the repeat > single contrast at session A. | 15 |
| Supplementary Table 11. Changes in activation between session A and D in all three groups (SI vs. MC, SW vs. MC) for switch and repeat trials in the ROIs defined on the switch > repeat contrast at session A. | 16 |
| Supplementary Table 12. Changes in activation across all sessions (A, B, C, and D) in the two practice groups (SI vs. SW) for switch and repeat trials in the ROIs defined on the switch > repeat contrast at session A. | 17 |
| Supplementary Results 1. Seed-to-voxel connectivity | 18 |
| Supplementary Results 2. IFJ to lPFC connectivity | 19 |
| Supplementary Results 3. Changes in Deviation Score and association with changes in connectivity | 20 |
| References | 22 |

Supplementary Table 1: Details of the training games children completed over nine weeks. Colors match the colors in Figure 1, such that the same colors indicate games with the same task structure. Opaque colors indicate one of the three repeating games and more translucent colors indicate one of the unique games.

| Type | Name | Task | Cues | Stimuli | Rule |
|---|---|---|---|---|---|
| Repeating | **Robot** | A robot is presented in the middle of the screen.<br><br>Sort the robots with a variable mapping of response keys depending on the spaceship cue | **Spaceships:**<br>Blue<br>Purple | **Robots:**<br>Robots (yellow-black eyes)<br>Robots (black-white eyes)<br>Robots (two different eye colors) | Focus on the robot's eyes |
| Unique | **Cat** | A cat is presented in the middle of the screen.<br><br>Sort the cats with a variable mapping of response keys depending on the food type | **Food types:**<br>Pouch food<br>Canned food | **Cats:**<br>Brown cats<br>Gray cats<br>Blue cats | Focus on the cat's color |
| Unique | **Socks** | Socks are presented in the middle of the screen.<br><br>Sort the socks with a variable mapping of response keys depending on the animal paws | **Animal paws:**<br>Dog paw<br>Bear paw | **Socks:**<br>Solid color socks<br>Striped socks<br>Socks with differently colored cuff | Focus on the socks pattern |
| Unique | **House** | A house is presented in the middle of the screen.<br><br>Sort the houses a variable the mapping of response keys depending on the key type | **Keys:**<br>Pretzel key<br>Ring key | **Houses:**<br>Stone houses<br>Boot houses<br>Mushroom houses | Focus on the type of house |
| Unique | **Vehicles** | A vehicle is presented in the middle of the screen.<br><br>Sort the vehicles with the mapping of response keys depending on the traffic light color | **Traffic light colors:**<br>Green<br>Red | **Vehicles:**<br>Cars<br>Motorcycles<br>Trucks | Focus on the type of vehicle |

| | | | | | |
|---|---|---|---|---|---|
| Repeating | **Monster** | Monsters as cue indicating which of the three tasks to perform<br><br>A shape in different colors and patterns are presented in the middle of the screen. | **Monsters:**<br>Curious<br>Fearful<br>Laughing | **Shapes:**<br>Circle<br>Square<br>Pentagon<br>**Patterns:**<br>Stripes<br>Wavy lines<br>Dots<br>**Colors:**<br>Yellow<br>Orange<br>Blue | **Shape Game:**<br>Focus on the shape<br>**Pattern Game:**<br>Focus on the pattern<br>**Color Game:**<br>Focus on the color |
| Unique | **Ice Cream** | Beach items as cue indicating which of the three tasks to perform<br><br>An ice cone with a varying shape, number of scoops, and scoop color presented in the middle of the screen. | **Beach items:**<br>Deck chair<br>Sunglasses<br>Beach umbrella | **Colors:**<br>Orange<br>Green<br>Pink<br>**Number of scoops:**<br>1<br>2<br>3<br>**Cone shape:**<br>Square<br>Cone<br>Pointed | **Color Game:**<br>Focus on the ice cream color<br>**Scoop Game:**<br>Focus on the number of scoops<br>**Cone Game:**<br>Focus on the cone shape |
| Unique | **Mandala** | Cartoon figures in different yoga poses as cue indicating which of the three tasks to perform<br><br>A mandala in different colors and patterns, with a gemstone in the center is presented in the middle of the screen. | **Yoga poses:**<br>Stretching<br>Seated<br>Standing | **Gemstones:**<br>Amber<br>Emerald<br>Sapphire<br>**Colors:**<br>Yellow<br>Green<br>Blue<br>**Shapes:**<br>Pointed<br>Round<br>Half pointed/half round | **Gemstone Game:**<br>Focus on the gemstone type in the center of the mandala<br>**Color Game:**<br>Focus on the mandala color<br>**Shape Game:**<br>Focus on the mandala shape |
| Unique | **Continent** | Animals as cue indicating which of the three tasks to perform<br><br>A continent in different colors with a circle around it is presented in the middle of the screen. | **Animals:**<br>Fox<br>Guinea pig<br>Squirrel | **Continent:**<br>Africa<br>Australia<br>Europe<br>**Color:**<br>Yellow<br>Green<br>Blue<br>**Circle:**<br>Short dashed<br>Long dashed<br>Solid | **Continent Game:**<br>Focus on the continent<br>**Color Game:**<br>Focus on the continent's color<br>**Circle Game:**<br>Focus on the circle border |
| Unique | **Cupcake** | Candle color as cue indicating which of the three tasks to perform.<br><br>A cupcake with | **Candle colors:**<br>Purple<br>Red<br>Green | **Toppings:**<br>Kiwi<br>Sprinkles<br>Gummy bears<br>**Icing colors:**<br>Green | **Topping Game:**<br>Focus on the topping<br>**Icing Game:**<br>Focus on the icing color |

| | | | | | |
|---|---|---|---|---|---|
| | | different toppings, icing color, and paper wrapper is presented in the middle of the screen. | | Brown<br>Purple<br>**Paper wrappers:**<br>Polka dot<br>Solid color<br>Striped | **Wrapper Game:** Focus on the wrapper pattern |
| Repeating | **Face-Scene** | A random combination of a face, a scene, and an object are presented simultaneously on the screen.<br><br>Geometric shapes as cue indicating which of the three tasks to perform | **Geometric shapes:**<br>Square<br>Circle<br>Hexagon | **Faces:**<br>Child<br>Young adult<br>Older adult<br>**Landscapes:**<br>Forest<br>Desert<br>Sea<br>**Objects:**<br>Yellow<br>Red<br>Purple | **Face Game:** Focus on the face<br>**Landscape Game:** Focus on the landscape<br>**Object Game:** Focus on the object's color |
| Unique | **Shelves** | A random combination of a toy, a book, and a pen are presented simultaneously on the screen.<br><br>Shelf colors as cue indicating which of the three tasks to perform | **Shelf colors:**<br>Brown<br>Green<br>Yellow | **Toys:**<br>Teddy bear<br>Horse<br>Block<br>**Books:**<br>Black<br>Red<br>Blue<br>**Pens:**<br>Pencil<br>Paintbrush<br>Ballpoint pen | **Toy Game:** Focus on the identity of the toy<br>**Book Game:** Focus on the book color<br>**Pen Game:** Focus on the pen |
| Unique | **Circus** | A random combination of a clown, a ball, and an animal are presented simultaneously on the screen.<br><br>Background color as cue indicating which of the three tasks to perform | **Background color:**<br>Red<br>Green<br>Blue | **Clown:**<br>Triangular hat<br>Flat hat<br>Jester hat<br>**Ball:**<br>Colorful<br>Patterned Striped<br>**Animal:**<br>Leopard<br>Lion<br>Tiger | **Clown Game:** Focus on the clown's hat<br>**Animal Game:** Focus on the animal<br>**Ball Game:** Focus on the ball's pattern |
| Unique | **Fields** | A random combination of a flower, a cloud, and a dog are presented simultaneously on the screen.<br><br>Background landscape as cue indicating which of the three tasks to perform | **Background landscape:**<br>Beach<br>Grass<br>Snow | **Flowers:**<br>Pink<br>Yellow<br>Purple<br>**Clouds:**<br>Rain cloud<br>Cloud with sun<br>Thundercloud<br>**Dogs:**<br>Bulldog<br>Pekingese<br>Dachshund dog | **Flower Game:** Focus on the flower color<br>**Cloud Game:** Focus on the cloud<br>**Dog Game:** Focus on the dog breed |
| Unique | **Sky** | A random | **Sky** | **Birds:** | **Bird Game:** |

| | | | | | |
|---|---|---|---|---|---|
| | | combination of a bird, fireworks, and an aircraft are presented simultaneously on the screen.<br><br>Sky background as cue indicating which of the three tasks to perform | **background:**<br>Blue sky<br>Night sky<br>Stormy sky | Parrot<br>Owl<br>Seagull<br>**Fireworks:**<br>Green<br>Red<br>Blue<br>**Aircraft:**<br>Balloon<br>Airplane<br>Helicopter | Focus on the bird type<br>**Firework Game:**<br>Focus on the firework color<br>**Aircraft Game:**<br>Focus on the aircraft |

Supplementary Table 2. Details of the distribution of trials in single and mixed blocks for the three types of training games for the high-intensity single-tasking (SI) and high-intensity task-switching (SW) groups.

| | SI group | | SW group | |
|---|---|---|---|---|
| Training task | single | mixed | single | mixed |
| Face-scene task and unique tasks of the same structure | 405 | 81 | 81 | 405 |
| Monster task and unique tasks of the same structure | 405 | 81 | 81 | 405 |
| Robot tasks and unique tasks of the same structure | 404 | 81 | 80 | 405 |

Supplementary Table 3a & 3b. Model comparisons showing that for behavioral measures the model including all interactions fit best or did not differ from best-fitting model. Models were compared using the LOO function within the loo package in R (Vehtari et al. 2022). ELPD difference indicates the difference between each model's Bayesian leave-one-out estimate of the expected log pointwise predictive density (ELPD) and the best fitting model, whole ELPD difference is zero. SE indicates the standard error of this difference. Note that there is not a fixed ratio of SE to ELPD difference to indicate significant differences between models. Until these exist, developers' suggestions are that an ELPD difference around five times the SE can be interpreted as indicating differences between models. Bold values indicate the best fitting model.

Supplementary Table 3a: Models across all three groups and sessions A and D.

| Model | Accuracy | | RT | |
|---|---|---|---|---|
| | ELPD difference | SE | ELPD difference | SE |
| condition x session x group | **0** | 0 | **0** | 0 |
| condition + session x group | 20.3 | 8.6 | 152.8 | 17.3 |
| condition x session + group | 11.9 | 6.6 | 12.5 | 4.5 |
| condition + session + group | 20 | 8.6 | 149.9 | 17.3 |

Supplementary Table 3b: Models for SI and SW groups and across all sessions A – D.

| Model | Accuracy | | RT | |
|---|---|---|---|---|
| | ELPD difference | SE | ELPD difference | SE |
| condition x session x group + condition x session(^2) x group | **0** | 0 | **0** | 0 |
| condition + session x group + session(^2) x group | 19.6 | 7.5 | 172.4 | 18.4 |
| condition x session + condition x session(^2) + group | 4.5 | 5 | 14.8 | 6.7 |
| condition + session + session(^2) + group | 25.6 | 7.5 | 173.1 | 18.5 |

Supplementary Table 4. Changes in activation across all sessions (A–D) in the two training groups (SI vs. SW) for repeat and single trials in the regions of interest (ROIs) defined on the repeat > single contrast in the adult comparison sample. CI indicates 95% credible intervals; bold values indicate estimates whose 95%-CI did not include zero. Dorsal anterior cingulate cortex (dACC), dorsolateral prefrontal cortex (dlPFC), inferior frontal junction (IFJ), superior parietal lobe (SPL)

| Effect | Left dACC | | | Left dlPFC | | | Left IFJ | | | Left SPL | | |
|---|---|---|---|---|---|---|---|---|---|---|---|---|
| | Estimate | CI | | Estimate | CI | | Estimate | CI | | Estimate | CI | |
| Intercept | **1.95** | 1.36 | 2.53 | **0.87** | 0.32 | 1.42 | **0.75** | 0.27 | 1.22 | **1.07** | 0.65 | 1.49 |
| group: SW vs. SI | -0.08 | -0.87 | 0.72 | 0.16 | -0.57 | 0.92 | 0.22 | -0.45 | 0.9 | 0.39 | -0.21 | 0.97 |
| session: B vs. A | -0.32 | -1.07 | 0.46 | 0.03 | -0.71 | 0.77 | 0.13 | -0.45 | 0.71 | 0.09 | -0.5 | 0.7 |
| session C vs. A | -0.32 | -1.06 | 0.41 | -0.26 | -0.98 | 0.49 | -0.33 | -0.91 | 0.25 | -0.01 | -0.58 | 0.58 |
| session D vs. A | -0.18 | -0.92 | 0.55 | -0.11 | -0.85 | 0.63 | -0.09 | -0.68 | 0.5 | 0.21 | -0.4 | 0.82 |
| condition: repeat vs. single | **1.53** | 0.84 | 2.24 | **1.69** | 0.94 | 2.45 | **1.28** | 0.65 | 1.91 | **1.57** | 0.95 | 2.18 |
| group SW x session B | 0.2 | -0.82 | 1.17 | -0.32 | -1.32 | 0.66 | -0.3 | -1.07 | 0.46 | -0.46 | -1.27 | 0.33 |
| group SW x session C | -0.21 | -1.18 | 0.77 | -0.41 | -1.42 | 0.58 | 0.07 | -0.73 | 0.83 | -0.24 | -1.04 | 0.55 |
| group SW x session D | -0.39 | -1.4 | 0.63 | -0.78 | -1.82 | 0.26 | -0.42 | -1.26 | 0.4 | -0.69 | -1.55 | 0.18 |
| group SW x repeat vs. single | 0.71 | -0.27 | 1.68 | -0.56 | -1.57 | 0.46 | -0.31 | -1.18 | 0.58 | 0.51 | -0.36 | 1.39 |
| session B x repeat vs. single | 0.44 | -0.63 | 1.51 | **-1.17** | -2.3 | -0.03 | -0.33 | -1.25 | 0.57 | -0.11 | -1.01 | 0.78 |
| session C x repeat vs. single | 0.72 | -0.31 | 1.78 | 0 | -1.13 | 1.09 | 0.3 | -0.59 | 1.17 | -0.46 | -1.38 | 0.43 |
| session D x repeat vs. single | -0.28 | -1.35 | 0.78 | -0.29 | -1.43 | 0.81 | 0.62 | -0.36 | 1.59 | -0.88 | -1.8 | 0.05 |
| group SW x session B x repeat | -0.74 | -2.16 | 0.7 | 0.72 | -0.79 | 2.22 | 0.25 | -0.99 | 1.47 | -0.14 | -1.38 | 1.1 |
| group SW x session C x repeat | -0.97 | -2.44 | 0.45 | -0.07 | -1.53 | 1.39 | -0.37 | -1.59 | 0.83 | -0.22 | -1.43 | 1.02 |
| group SW x session D x repeat | -0.07 | -1.53 | 1.39 | 0.6 | -0.95 | 2.19 | 0.24 | -1.12 | 1.58 | 0.06 | -1.25 | 1.4 |

| Effect | Right dACC | | | Right dlPFC | | | Right IFJ | | | Right SPL | | |
|---|---|---|---|---|---|---|---|---|---|---|---|---|
| | Estimate | CI | | Estimate | CI | | Estimate | CI | | Estimate | CI | |
| Intercept | **1.41** | 0.94 | 1.9 | **0.68** | 0.13 | 1.22 | **0.65** | 0.21 | 1.1 | **0.75** | 0.2 | 1.31 |
| group: SW vs. SI | -0.12 | -0.79 | 0.58 | -0.12 | -0.88 | 0.63 | -0.41 | -1.05 | 0.23 | 0.09 | -0.68 | 0.85 |
| session: B vs. A | -0.29 | -0.93 | 0.37 | 0.15 | -0.59 | 0.88 | 0.25 | -0.34 | 0.83 | 0.44 | -0.29 | 1.15 |
| session C vs. A | -0.26 | -0.93 | 0.41 | -0.36 | -1.17 | 0.44 | -0.28 | -0.85 | 0.32 | 0.03 | -0.7 | 0.76 |
| session D vs. A | -0.29 | -0.97 | 0.4 | 0.1 | -0.63 | 0.85 | 0.07 | -0.51 | 0.64 | 0.17 | -0.58 | 0.91 |
| condition: repeat vs. single | **1.33** | 0.71 | 1.96 | **1.04** | 0.28 | 1.82 | **1.36** | 0.78 | 1.96 | **1.69** | 0.95 | 2.42 |
| group SW x session B | 0.27 | -0.6 | 1.13 | -0.16 | -1.17 | 0.83 | -0.03 | -0.82 | 0.75 | -0.57 | -1.51 | 0.4 |
| group SW x session C | 0.14 | -0.75 | 1.04 | 0.18 | -0.9 | 1.26 | 0.45 | -0.37 | 1.24 | -0.18 | -1.15 | 0.77 |
| group SW x session D | -0.05 | -1.02 | 0.91 | -0.63 | -1.71 | 0.43 | -0.28 | -1.11 | 0.54 | -0.44 | -1.47 | 0.61 |
| group SW x repeat vs. single | 0.49 | -0.4 | 1.37 | -0.25 | -1.3 | 0.8 | -0.45 | -1.29 | 0.38 | -0.41 | -1.39 | 0.58 |
| session B x repeat vs. single | 0.48 | -0.48 | 1.43 | -0.51 | -1.64 | 0.63 | -0.6 | -1.47 | 0.28 | -0.45 | -1.56 | 0.66 |
| session C x repeat vs. single | 0.22 | -0.72 | 1.15 | 0.06 | -1.1 | 1.21 | -0.13 | -1.03 | 0.75 | -0.75 | -1.87 | 0.36 |
| session D x repeat vs. single | -0.47 | -1.44 | 0.48 | -0.7 | -1.8 | 0.37 | -0.78 | -1.65 | 0.07 | **-1.35** | -2.45 | -0.25 |
| group SW x session B x repeat | -0.82 | -2.09 | 0.47 | 0.23 | -1.27 | 1.73 | 0.36 | -0.85 | 1.54 | 0.25 | -1.18 | 1.65 |
| group SW x session C x repeat | -0.88 | -2.15 | 0.41 | -0.78 | -2.31 | 0.75 | -0.52 | -1.72 | 0.65 | 0.18 | -1.28 | 1.65 |
| group SW x session D x repeat | -0.26 | -1.6 | 1.11 | 0.25 | -1.32 | 1.79 | 0.64 | -0.57 | 1.83 | 0.49 | -1 | 1.99 |

Supplementary Table 5: Changes in activation across all sessions (A – D) in the two training groups (SI vs. SW) for switch and repeat trials in the ROIs defined on the switch > repeat contrast in the adult comparison sample. CI denotes 95%-Credible Intervals; bold values indicate estimates whose 95%-CI did not include zero. Supplementary motor area (SMA)

| Effect | Left dACC | | | Left IFJ | | | Left Precuneus | | | Left SMA | | | Left SPL | | | Right dACC | | | Right Precuneus | | | Right SMA | | |
|---|---|---|---|---|---|---|---|---|---|---|---|---|---|---|---|---|---|---|---|---|---|---|---|---|
| | Estimate | CI | | Estimate | CI | | Estimate | CI | | Estimate | CI | | Estimate | CI | | Estimate | CI | | Estimate | CI | | Estimate | CI | |
| Intercept | **2.45** | 1.69 | 3.19 | **2.65** | 1.81 | 3.52 | -0.46 | -1.14 | 0.23 | **3.15** | 2.34 | 3.97 | **2.17** | 1.54 | 2.8 | **2.15** | 1.54 | 2.78 | 0.25 | -0.49 | 0.97 | **2.46** | 1.7 | 3.23 |
| group: SW vs. SI | 0.07 | -0.96 | 1.06 | -0.22 | -1.41 | 0.96 | 0.07 | -0.89 | 1.01 | 0.71 | -0.43 | 1.84 | 0.38 | -0.5 | 1.27 | -0.24 | -1.09 | 0.61 | 0.24 | -0.74 | 1.24 | 0.33 | -0.7 | 1.36 |
| session: B vs. A | 0.09 | -0.82 | 0.98 | -0.7 | -1.78 | 0.37 | 0.72 | -0.09 | 1.53 | -0.28 | -1.29 | 0.74 | -0.16 | -0.91 | 0.56 | 0.11 | -0.78 | 1 | **1.24** | 0.42 | 2.05 | 0.1 | -0.84 | 1.04 |
| session C vs. A | 0.56 | -0.45 | 1.59 | -0.25 | -1.23 | 0.7 | 0.58 | -0.28 | 1.43 | 0.3 | -0.72 | 1.33 | -0.17 | -1.05 | 0.72 | 0.34 | -0.63 | 1.27 | 0.73 | -0.2 | 1.66 | 0.33 | -0.74 | 1.41 |
| session D vs. A | -0.53 | -1.55 | 0.51 | -0.08 | -1.14 | 0.95 | 0.52 | -0.46 | 1.46 | -0.58 | -1.84 | 0.62 | **-0.93** | -1.76 | -0.09 | -0.49 | -1.43 | 0.42 | 0.36 | -0.73 | 1.4 | -0.19 | -1.26 | 0.91 |
| condition: switch vs. repeat | 0.43 | -0.13 | 0.99 | **0.67** | 0.17 | 1.21 | **0.75** | 0.11 | 1.38 | 0.46 | -0.09 | 1.02 | **0.83** | 0.34 | 1.34 | 0.48 | -0.1 | 1.08 | 0.48 | -0.2 | 1.21 | 0.44 | -0.16 | 1.05 |
| group SW x session B | 0.01 | -1.19 | 1.21 | 0.71 | -0.69 | 2.1 | 0.32 | -0.78 | 1.44 | 0.05 | -1.3 | 1.4 | -0.11 | -1.12 | 0.89 | 0.13 | -1.05 | 1.34 | -0.37 | -1.46 | 0.71 | -0.26 | -1.51 | 1.01 |
| group SW x session C | -0.8 | -2.18 | 0.56 | -0.11 | -1.37 | 1.2 | -0.18 | -1.34 | 0.99 | -0.58 | -1.96 | 0.78 | -0.31 | -1.53 | 0.87 | -0.4 | -1.65 | 0.91 | -0.36 | -1.59 | 0.88 | -0.21 | -1.66 | 1.23 |
| group SW x session D | 0.26 | -1.16 | 1.67 | -0.22 | -1.69 | 1.24 | -0.09 | -1.42 | 1.28 | 0.02 | -1.7 | 1.74 | -0.03 | -1.2 | 1.16 | 0.2 | -1.1 | 1.48 | -0.12 | -1.57 | 1.37 | -0.02 | -1.57 | 1.49 |
| group SW x single | 0.06 | -0.73 | 0.83 | -0.09 | -0.81 | 0.59 | -0.07 | -0.98 | 0.82 | -0.06 | -0.84 | 0.73 | -0.35 | -1.07 | 0.36 | -0.09 | -0.92 | 0.73 | -0.08 | -1.02 | 0.84 | -0.09 | -0.94 | 0.74 |
| session B x switch | -0.47 | -1.3 | 0.36 | -0.71 | -1.51 | 0.03 | -0.3 | -1.28 | 0.65 | -0.53 | -1.36 | 0.28 | **-0.99** | -1.76 | -0.23 | -0.52 | -1.41 | 0.36 | 0.1 | -0.93 | 1.1 | -0.49 | -1.38 | 0.39 |
| session C x switch | -0.22 | -1.03 | 0.59 | -0.52 | -1.27 | 0.21 | -0.03 | -0.98 | 0.92 | -0.16 | -0.97 | 0.64 | -0.67 | -1.39 | 0.05 | -0.3 | -1.18 | 0.57 | 0.01 | -0.97 | 1.01 | -0.26 | -1.16 | 0.62 |
| session D x switch | 0.45 | -0.4 | 1.3 | -0.14 | -0.89 | 0.61 | -0.21 | -1.22 | 0.77 | -0.39 | -1.23 | 0.43 | -0.47 | -1.23 | 0.28 | 0.29 | -0.6 | 1.19 | 0.08 | -0.97 | 1.13 | -0.62 | -1.53 | 0.29 |
| group SW x session B x switch | 0.45 | -0.66 | 1.55 | 0.39 | -0.61 | 1.43 | -0.24 | -1.53 | 1.05 | 0.6 | -0.5 | 1.71 | 0.77 | -0.25 | 1.83 | 0.67 | -0.51 | 1.84 | -0.35 | -1.66 | 0.99 | 0.57 | -0.58 | 1.8 |
| group SW x session C x switch | 0.32 | -0.78 | 1.42 | 0.51 | -0.47 | 1.51 | 0.04 | -1.25 | 1.32 | 0.19 | -0.93 | 1.31 | 0.8 | -0.21 | 1.8 | 0.55 | -0.64 | 1.73 | 0.15 | -1.15 | 1.48 | 0.15 | -1.06 | 1.39 |
| group SW x session D x switch | -0.66 | -1.84 | 0.51 | -0.04 | -1.04 | 0.99 | -0.57 | -1.97 | 0.8 | 0.41 | -0.75 | 1.58 | 0.19 | -0.88 | 1.28 | -0.46 | -1.74 | 0.83 | -0.28 | -1.7 | 1.13 | 0.47 | -0.79 | 1.76 |

Supplementary Table 6. Details on ROIs defined on the adult comparison sample

| Contrast | Hemisphere | ROI | Size | Peak | | |
|---|---|---|---|---|---|---|
| Repeat > single | left | dlPFC | 243 | -44 | 24 | 28 |
| | | IFJ | 381 | -42 | 4 | 32 |
| | | dACC | 122 | -4 | 12 | 52 |
| | | SPL | 332 | -28 | -58 | 48 |
| | right | dlPFC | 239 | 48 | 30 | 26 |
| | | IFJ | 410 | 38 | 0 | 40 |
| | | SPL | 249 | 32 | -52 | 50 |
| | | dACC | 107 | 8 | 12 | 52 |
| Switch > repeat | left | dACC | 218 | -6 | 10 | 50 |
| | | SMA | 407 | -6 | 8 | 52 |
| | | IFJ | 102 | -48 | 2 | 32 |
| | | SPL | 509 | -32 | -52 | 58 |
| | | Precuneus | 569 | -10 | -72 | 40 |
| | right | dACC | 123 | 12 | 20 | 36 |
| | | SMA | 216 | 2 | 2 | 56 |
| | | Precuneus | 451 | 20 | -58 | 6 |

Supplementary Table 7. Model outputs of performance (accuracy and response times) between sessions A and D in the two training groups compared to the passive control group (i.e., SI vs. PC, SW vs. PC) for single, repeat, and switch trials (repeat as reference level). CI denotes 95%-Credible Intervals; bold values indicate estimates whose 95%-CI did not include zero.

| Effect | Accuracy | | | RT | | |
|---|---|---|---|---|---|---|
| | Estimate | CI | | Estimate | CI | |
| Intercept | **1.77** | 1.39 | 2.15 | **1.46** | 1.41 | 1.52 |
| condition: single vs. repeat | **1.65** | 1.35 | 1.96 | **-0.24** | -0.28 | -0.2 |
| condition: switch vs. repeat | **-0.51** | -0.67 | -0.35 | **0.2** | 0.17 | 0.23 |
| session: D vs. A | -0.14 | -0.55 | 0.26 | -0.01 | -0.08 | 0.05 |
| group: SI vs. PC | -0.12 | -0.56 | 0.32 | 0.01 | -0.06 | 0.07 |
| group: SW vs. PC | 0.13 | -0.31 | 0.57 | 0.01 | -0.05 | 0.08 |
| condition (single vs. repeat) x session (D vs. A) | **-0.7** | -0.97 | -0.44 | 0.03 | 0 | 0.06 |
| condition (switch vs. repeat) x session (D vs. A) | 0.16 | -0.03 | 0.36 | **-0.06** | -0.09 | -0.03 |
| condition (single vs. repeat) x group (SI vs. PC) | **-0.54** | -0.88 | -0.19 | 0.02 | -0.03 | 0.07 |
| condition (switch vs. repeat) x group (SI vs. PC) | -0.1 | -0.29 | 0.08 | -0.02 | -0.06 | 0.02 |
| condition (single vs. repeat) x group (SW vs. PC) | **-0.43** | -0.78 | -0.08 | 0.01 | -0.04 | 0.06 |
| condition (switch vs. repeat) x group (SW vs. PC) | -0.16 | -0.35 | 0.03 | 0 | -0.03 | 0.04 |
| session (D vs. A) x group (SI vs. PC) | 0.05 | -0.42 | 0.5 | **-0.09** | -0.16 | -0.02 |
| session (D vs. A) x group (SW vs. PC) | **0.61** | 0.13 | 1.09 | **-0.09** | -0.16 | -0.01 |
| condition (single vs. repeat) x group (SI vs. PC) x session (D vs. A) | **0.98** | 0.66 | 1.3 | **0.06** | 0.02 | 0.09 |
| condition (switch vs. repeat) x group (SI vs. PC) x session (D vs. A) | 0.07 | -0.17 | 0.31 | 0 | -0.04 | 0.03 |
| condition (single vs. repeat) x group (SW vs. PC) x session (D vs. A) | **0.48** | 0.12 | 0.84 | 0.04 | 0 | 0.08 |
| condition (switch vs. repeat) x group (SW vs. PC) x session (D vs. A | 0.05 | -0.22 | 0.31 | 0.02 | -0.01 | 0.06 |

Supplementary Table 8. Model outputs of performance (accuracy and response times) across all sessions (A – D) in the two training groups (SW vs. SI) for single, repeat, and switch trials (repeat as reference level). CI denotes 95%-Credible Intervals; bold values indicate estimates whose 95%-CI did not include zero.

| Effect | Accuracy | | | RT | | |
|---|---|---|---|---|---|---|
| | Estimate | CI | | Estimate | CI | |
| Intercept | **1.62** | 1.32 | 1.93 | **1.47** | 1.43 | 1.52 |
| session | **0.62** | 0.46 | 0.78 | **-0.07** | -0.09 | -0.05 |
| group: SW vs. SI | 0.31 | -0.07 | 0.7 | 0.04 | -0.02 | 0.09 |
| condition: single vs. repeat | **0.89** | 0.72 | 1.06 | **-0.23** | -0.26 | -0.2 |
| condition: switch vs. repeat | **-0.57** | -0.68 | -0.47 | **0.17** | 0.15 | 0.19 |
| session^2 | **-0.22** | -0.26 | -0.18 | **0.02** | 0.01 | 0.02 |
| session x group (SW vs. SI) | 0.14 | -0.09 | 0.38 | -0.02 | -0.06 | 0.01 |
| session x condition (single vs. repeat) | **0.5** | 0.29 | 0.72 | 0.02 | 0 | 0.04 |
| session x condition (switch vs. repeat) | -0.07 | -0.23 | 0.09 | **0.03** | 0.01 | 0.05 |
| condition (single vs. repeat) x group (SW vs. SI) | 0.11 | -0.12 | 0.34 | -0.03 | -0.06 | 0.01 |
| condition (switch vs. repeat) x group (SW vs. SI) | -0.08 | -0.23 | 0.06 | 0.03 | 0 | 0.06 |
| session^2 x group (SW vs. SI) | -0.02 | -0.08 | 0.04 | 0 | 0 | 0.01 |
| session^2 x condition (single vs. repeat) | **-0.13** | -0.2 | -0.06 | 0 | -0.01 | 0.01 |
| session^2 x condition (switch vs. repeat) | 0.04 | -0.01 | 0.09 | **-0.02** | -0.02 | -0.01 |
| session x condition (single vs. repeat) x group (SW vs. SI) | **-0.48** | -0.8 | -0.16 | **0.05** | 0.02 | 0.08 |
| session x condition (switch vs. repeat) x group (SW vs. SI) | 0.1 | -0.14 | 0.34 | **-0.05** | -0.08 | -0.02 |
| session^2 x condition (single vs. repeat) x group (SW vs. SI) | 0.11 | 0 | 0.22 | **-0.02** | -0.03 | -0.01 |
| session^2 x condition (switch vs. repeat) x group (SW vs. SI) | -0.03 | -0.11 | 0.05 | **0.02** | 0.01 | 0.03 |

Supplementary Table 9. Changes in activation between session A and D in all three groups (SI vs. PC [passive control group], SW vs. PC) for repeat and single trials in the ROIs defined on the repeat > single contrast at session A. CI denotes 95%-Credible Intervals; bold values indicate estimates whose 95%-CI did not include zero.

| Effect | Left IFJ | | | Left SPL | | | left dlPFC | | | Right IFJ | | | Right SPL | | |
|---|---|---|---|---|---|---|---|---|---|---|---|---|---|---|---|
| | Estimate | CI | | Estimate | CI | | Estimate | CI | | Estimate | CI | | Estimate | CI | |
| Intercept | 0.62 | -0.01 | 1.23 | **1.12** | 0.52 | 1.7 | 0.28 | -0.38 | 0.93 | 0.5 | -0.06 | 1.04 | **0.73** | 0.14 | 1.31 |
| group: SI vs. PC | 0.14 | -0.7 | 0.97 | 0.02 | -0.77 | 0.81 | 0.43 | -0.42 | 1.3 | 0.09 | -0.65 | 0.83 | 0.05 | -0.72 | 0.83 |
| group: SW vs. PC | 0.32 | -0.49 | 1.15 | 0.4 | -0.37 | 1.18 | 0.68 | -0.15 | 1.54 | -0.26 | -0.98 | 0.48 | 0.11 | -0.66 | 0.87 |
| session: D vs. A | 0.38 | -0.26 | 1.05 | 0.27 | -0.5 | 1.05 | 0.56 | -0.26 | 1.36 | 0.19 | -0.46 | 0.82 | 0.21 | -0.53 | 0.93 |
| condition: repeat vs. single | **2.34** | 1.7 | 2.98 | **3.09** | 2.29 | 3.87 | **2.05** | 1.24 | 2.88 | **2.13** | 1.47 | 2.79 | **2.49** | 1.75 | 3.24 |
| groupSI x session D | -0.42 | -1.33 | 0.49 | -0.07 | -1.15 | 1 | -0.54 | -1.7 | 0.55 | -0.08 | -0.96 | 0.79 | 0.04 | -0.98 | 1.06 |
| groupSW x session D | -0.87 | -1.78 | 0.02 | -0.72 | -1.77 | 0.35 | **-1.41** | -2.52 | -0.28 | -0.44 | -1.34 | 0.44 | -0.44 | -1.44 | 0.58 |
| group SI x repeat | **-0.96** | -1.83 | -0.1 | **-1.33** | -2.39 | -0.3 | -0.29 | -1.39 | 0.82 | -0.83 | -1.7 | 0.05 | -0.71 | -1.75 | 0.32 |
| group SW x repeat | **-1.36** | -2.23 | -0.51 | -0.73 | -1.78 | 0.32 | -0.94 | -2.03 | 0.12 | **-1.24** | -2.12 | -0.41 | -0.92 | -1.91 | 0.05 |
| session D x repeat | -0.33 | -1.25 | 0.58 | -0.59 | -1.73 | 0.54 | -0.96 | -2.18 | 0.21 | -0.75 | -1.67 | 0.18 | -0.8 | -1.87 | 0.28 |
| group SI x session D x repeat | 0.56 | -0.7 | 1.83 | -0.32 | -1.85 | 1.23 | 0.48 | -1.13 | 2.14 | 0.07 | -1.21 | 1.33 | -0.26 | -1.83 | 1.26 |
| groupSW x session D x repeat | 1.03 | -0.22 | 2.3 | -0.36 | -1.91 | 1.2 | 1.11 | -0.48 | 2.74 | 0.67 | -0.58 | 1.92 | -0.1 | -1.59 | 1.35 |

Supplementary Table 10. Changes in activation across all sessions (A, B, C, and D) in the two practice groups (SI vs. SW) for repeat and single trials in the ROIs defined on the repeat > single contrast at session A. CI denotes 95%-Credible Intervals; bold values indicate estimates whose 95%-CI did not include zero.

| Effect | Left IFJ | | | Left SPL | | | left dlPFC | | | Right IFJ | | | Right SPL | | |
|---|---|---|---|---|---|---|---|---|---|---|---|---|---|---|---|
| | Estimate | CI | | Estimate | CI | | Estimate | CI | | Estimate | CI | | Estimate | CI | |
| Intercept | **0.78** | 0.35 | 1.22 | **1.16** | 0.7 | 1.62 | **0.72** | 0.17 | 1.27 | **0.6** | 0.16 | 1.04 | **0.8** | 0.28 | 1.31 |
| group: SW vs. SI | 0.17 | -0.46 | 0.79 | 0.35 | -0.29 | 1 | 0.24 | -0.51 | 1.01 | -0.36 | -0.96 | 0.27 | 0.06 | -0.65 | 0.77 |
| session: B vs. A | 0.09 | -0.42 | 0.6 | -0.01 | -0.63 | 0.62 | 0.08 | -0.68 | 0.84 | 0.26 | -0.31 | 0.84 | 0.42 | -0.24 | 1.07 |
| session C vs. A | -0.31 | -0.79 | 0.19 | -0.05 | -0.65 | 0.58 | -0.31 | -1.04 | 0.45 | -0.25 | -0.81 | 0.36 | 0.06 | -0.6 | 0.73 |
| session D vs. A | -0.06 | -0.56 | 0.44 | 0.18 | -0.47 | 0.83 | -0.04 | -0.78 | 0.7 | 0.09 | -0.49 | 0.66 | 0.18 | -0.51 | 0.87 |
| condition: repeat vs. single | **1.23** | 0.68 | 1.77 | **1.68** | 1.04 | 2.33 | **1.74** | 0.99 | 2.48 | **1.32** | 0.75 | 1.9 | **1.84** | 1.13 | 2.56 |
| group SW x session B | -0.27 | -0.94 | 0.41 | -0.35 | -1.2 | 0.48 | -0.3 | -1.31 | 0.71 | -0.04 | -0.81 | 0.73 | -0.55 | -1.41 | 0.31 |
| group SW x session C | 0.04 | -0.65 | 0.73 | -0.18 | -1.02 | 0.68 | -0.3 | -1.31 | 0.68 | 0.41 | -0.38 | 1.2 | -0.19 | -1.09 | 0.71 |
| group SW x session D | -0.46 | -1.18 | 0.25 | -0.67 | -1.58 | 0.26 | -0.83 | -1.86 | 0.21 | -0.33 | -1.16 | 0.49 | -0.4 | -1.34 | 0.53 |
| group SW x repeat | -0.37 | -1.15 | 0.41 | 0.69 | -0.22 | 1.62 | -0.63 | -1.62 | 0.4 | -0.43 | -1.23 | 0.37 | -0.33 | -1.27 | 0.58 |
| session B x repeat | -0.43 | -1.24 | 0.38 | 0.05 | -0.87 | 0.97 | **-1.38** | -2.51 | -0.23 | -0.59 | -1.46 | 0.26 | -0.3 | -1.33 | 0.73 |
| session C x repeat | 0.29 | -0.48 | 1.06 | -0.37 | -1.33 | 0.57 | -0.09 | -1.18 | 1.03 | -0.12 | -0.97 | 0.72 | -0.72 | -1.78 | 0.3 |
| session D x repeat | 0.37 | -0.44 | 1.17 | -0.78 | -1.77 | 0.21 | -0.48 | -1.6 | 0.63 | -0.71 | -1.57 | 0.14 | **-1.06** | -2.12 | -0.02 |
| group SW x session B x repeat | 0.32 | -0.79 | 1.42 | -0.34 | -1.64 | 0.94 | 0.75 | -0.79 | 2.23 | 0.33 | -0.82 | 1.49 | 0.11 | -1.23 | 1.45 |
| group SW x session C x repeat | -0.24 | -1.31 | 0.84 | -0.5 | -1.79 | 0.81 | -0.15 | -1.66 | 1.33 | -0.51 | -1.68 | 0.64 | 0.18 | -1.18 | 1.49 |
| group SW x session D x repeat | 0.48 | -0.66 | 1.66 | -0.12 | -1.53 | 1.26 | 0.64 | -0.89 | 2.14 | 0.59 | -0.61 | 1.78 | 0.22 | -1.16 | 1.6 |

Supplementary Table 11. Changes in activation between session A and D in all three groups (SI vs. PC, SW vs. PC) for switch and repeat trials in the ROIs defined on the switch > repeat contrast at session A. CI denotes 95%-Credible Intervals; bold values indicate estimates whose 95%-CI did not include zero.

| Effect | Left IFJ | | | Left SPL | | | left Precuneus | | | Right Precuneus | | | Right SPL | | |
|---|---|---|---|---|---|---|---|---|---|---|---|---|---|---|---|
| | Estimate | CI | | Estimate | CI | | Estimate | CI | | Estimate | CI | | Estimate | CI | |
| Intercept | **3.67** | 2.57 | 4.79 | **3.67** | 2.92 | 4.41 | 0.39 | -0.39 | 1.16 | **1.73** | 0.83 | 2.65 | 0.61 | -0.09 | 1.34 |
| group: SI vs. PC | -0.89 | -2.39 | 0.64 | -0.93 | -1.93 | 0.09 | **-1.26** | -2.3 | -0.21 | **-1.97** | -3.21 | -0.74 | -0.47 | -1.44 | 0.52 |
| group: SW vs. PC | -1.2 | -2.67 | 0.32 | -0.39 | -1.37 | 0.59 | **-1.05** | -2.08 | -0.01 | **-1.53** | -2.74 | -0.33 | -0.86 | -1.79 | 0.08 |
| session: D vs. A | 0.05 | -1.08 | 1.16 | -0.49 | -1.42 | 0.45 | -0.6 | -1.54 | 0.33 | -0.75 | -1.78 | 0.32 | **-0.76** | -1.55 | -0.01 |
| condition: switch vs. repeat | 0.15 | -0.46 | 0.75 | 0.01 | -0.59 | 0.59 | 0.2 | -0.51 | 0.92 | 0.36 | -0.44 | 1.15 | 0.17 | -0.34 | 0.69 |
| groupSI x session D | -0.36 | -1.85 | 1.13 | -0.17 | -1.43 | 1.09 | **1.49** | 0.16 | 2.79 | 1.32 | -0.12 | 2.78 | 0.31 | -0.68 | 1.36 |
| groupSW x session D | -0.05 | -1.57 | 1.52 | -0.46 | -1.73 | 0.82 | **1.61** | 0.32 | 2.95 | 1.05 | -0.4 | 2.49 | 0.41 | -0.63 | 1.48 |
| group SI x switch | 0.63 | -0.17 | 1.45 | **1** | 0.2 | 1.8 | 0.45 | -0.53 | 1.42 | 0.19 | -0.87 | 1.28 | 0.33 | -0.33 | 0.99 |
| group SW x switch | 0.59 | -0.16 | 1.36 | 0.55 | -0.22 | 1.34 | 0.7 | -0.24 | 1.66 | -0.03 | -1.08 | 1.03 | 0.09 | -0.56 | 0.74 |
| session D x switch | 0.48 | -0.32 | 1.31 | -0.01 | -0.85 | 0.83 | 0.42 | -0.6 | 1.42 | 0.03 | -1.1 | 1.15 | -0.22 | -0.94 | 0.47 |
| group SI x session D x switch | -0.73 | -1.88 | 0.38 | -0.63 | -1.79 | 0.53 | -0.43 | -1.88 | 1.03 | 0 | -1.53 | 1.57 | -0.48 | -1.42 | 0.51 |
| groupSW x session D x switch | -0.87 | -2 | 0.25 | -0.25 | -1.39 | 0.9 | **-1.6** | -3.01 | -0.19 | -0.2 | -1.76 | 1.35 | -0.1 | -1.06 | 0.85 |

Supplementary Table 12: Changes in activation across all sessions (A, B, C, and D) in the two practice groups (SI vs. SW) for switch and repeat trials in the ROIs defined on the switch > repeat contrast at session A. CI denotes 95%-Credible Intervals.

| Effect | Left IFJ | | | Left SPL | | | left Precuneus | | | Right Precuneus | | | Right SPL | | |
|---|---|---|---|---|---|---|---|---|---|---|---|---|---|---|---|
| | Estimate | CI | | Estimate | CI | | Estimate | CI | | Estimate | CI | | Estimate | CI | |
| Intercept | **2.81** | 1.85 | 3.78 | **2.74** | 2.08 | 3.4 | **-0.85** | -1.51 | -0.16 | -0.04 | -0.82 | 0.72 | 0.22 | -0.32 | 0.77 |
| group: SW vs. SI | -0.3 | -1.63 | 1.05 | 0.51 | -0.38 | 1.38 | 0.17 | -0.75 | 1.1 | 0.22 | -0.81 | 1.25 | -0.5 | -1.29 | 0.26 |
| session: B vs. A | -0.82 | -1.86 | 0.24 | -0.17 | -0.96 | 0.59 | **1.01** | 0.19 | 1.83 | **1.24** | 0.41 | 2.07 | -0.23 | -0.91 | 0.44 |
| session C vs. A | -0.42 | -1.41 | 0.53 | -0.3 | -1.19 | 0.6 | 0.6 | -0.27 | 1.47 | 0.74 | -0.17 | 1.65 | -0.21 | -0.94 | 0.51 |
| session D vs. A | -0.32 | -1.38 | 0.69 | -0.73 | -1.57 | 0.11 | 0.91 | -0.08 | 1.86 | 0.44 | -0.62 | 1.48 | -0.51 | -1.16 | 0.16 |
| condition: switch vs. repeat | **0.73** | 0.26 | 1.21 | **0.96** | 0.46 | 1.46 | 0.63 | -0.01 | 1.26 | 0.56 | -0.11 | 1.25 | **0.48** | 0.08 | 0.89 |
| group SW x session B | 1.02 | -0.4 | 2.39 | -0.02 | -1.04 | 1.04 | 0.24 | -0.85 | 1.35 | -0.47 | -1.58 | 0.61 | 0.43 | -0.46 | 1.3 |
| group SW x session C | 0.16 | -1.13 | 1.47 | -0.32 | -1.53 | 0.94 | 0.18 | -0.96 | 1.36 | -0.26 | -1.45 | 0.97 | 0.3 | -0.66 | 1.23 |
| group SW x session D | 0.19 | -1.27 | 1.67 | -0.3 | -1.48 | 0.86 | 0.09 | -1.28 | 1.45 | -0.17 | -1.63 | 1.29 | 0.23 | -0.69 | 1.15 |
| group SW x switch | 0 | -0.62 | 0.64 | -0.39 | -1.09 | 0.32 | 0.28 | -0.58 | 1.16 | -0.17 | -1.1 | 0.76 | -0.22 | -0.78 | 0.34 |
| session B x switch | -0.67 | -1.37 | 0.02 | **-1.06** | -1.81 | -0.3 | 0.01 | -0.93 | 0.93 | 0.14 | -0.86 | 1.14 | -0.47 | -1.09 | 0.15 |
| session C x switch | -0.21 | -0.91 | 0.45 | -0.56 | -1.3 | 0.18 | 0.35 | -0.59 | 1.28 | -0.03 | -1.02 | 0.93 | **-0.68** | -1.3 | -0.09 |
| session D x switch | -0.21 | -0.91 | 0.5 | -0.6 | -1.34 | 0.16 | -0.02 | -0.99 | 0.94 | 0.04 | -1 | 1.04 | **-0.66** | -1.32 | -0.01 |
| group SW x session B x switch | 0.36 | -0.57 | 1.3 | 0.82 | -0.18 | 1.86 | -0.8 | -2.04 | 0.47 | -0.21 | -1.55 | 1.13 | 0.17 | -0.63 | 0.99 |
| group SW x session C x switch | -0.05 | -0.97 | 0.86 | 0.74 | -0.26 | 1.75 | -0.66 | -1.92 | 0.61 | 0.27 | -1.06 | 1.59 | 0.49 | -0.31 | 1.32 |
| group SW x session D x switch | -0.16 | -1.15 | 0.83 | 0.33 | -0.73 | 1.36 | -1.12 | -2.45 | 0.19 | -0.19 | -1.58 | 1.24 | 0.32 | -0.58 | 1.2 |

**Supplementary Results 1: Seed-to-voxel connectivity**

*Connectivity associated with mixing demand*
Whole-brain seed-to-voxel analyses with seeds in the left IFJ and SPL for repeat > single did not reveal any clusters that showed changes in connectivity with training (p < .05 FDR-corrected, voxel threshold at p < .001 uncorrected). As noted in the preregistration, we additionally conducted seed-to-voxel analyses for regions that showed changes in activation with training. More specifically, we analyzed whole-brain connectivity for a seed in the left dlPFC. No clusters showed changes in connectivity with the dlPFC with training (p < .05 FDR-corrected, voxel threshold at p < .001 uncorrected).

*Connectivity associated with switch demand*
The whole-brain seed-to-voxel analysis with a seed in the left IFJ revealed a cluster in the left angular gyrus that showed changes in connectivity for switch > repeat with training (p < .05 FDR-corrected, voxel threshold at p < .001 uncorrected), such that condition differences increased from session A to C in the SI group and then decreased again at session D, while condition differences in the SW group only increased at session D (see Figure SR1.1). There was no significant cluster for the left SPL seed region.

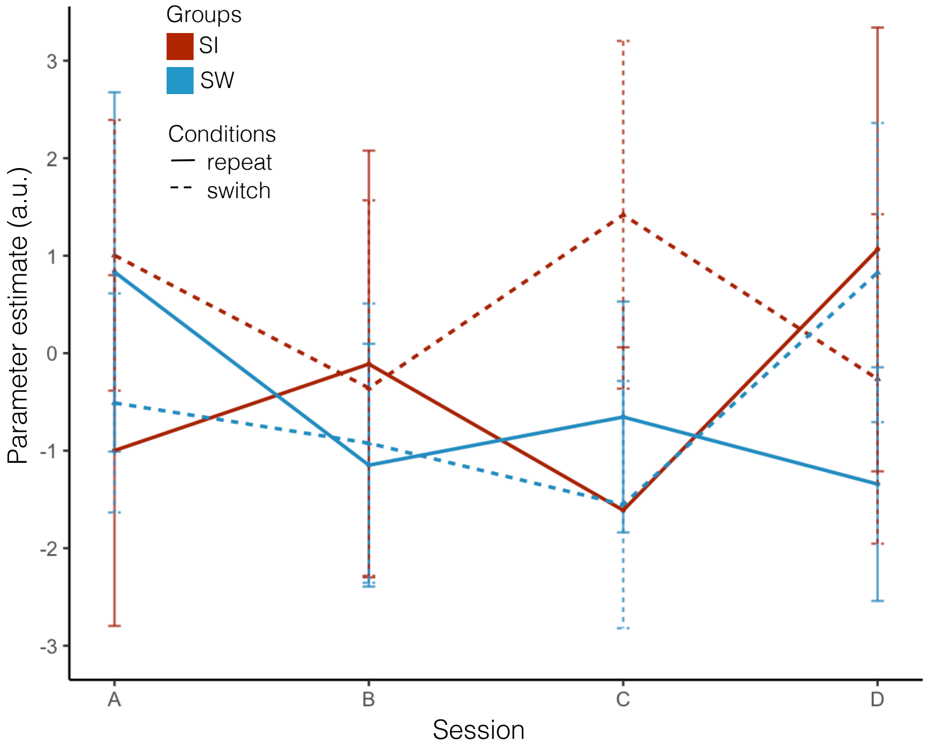


*Figure SR1.1: Change in connectivity between the IFJ and a cluster in the left angular gyrus (AG).* PPI parameters for the connection between the seed in the IFJ and the cluster in the left AG that showed change in the connectivity difference between repeat and switch trials. Error bars denote 95%-confidence intervals. The single-task (SI) practice group is shown in red and the task-switching practice group in blue. Connectivity for repeat trials is shown in the solid line and for switch trials in the long-dashed line for each session.

### Supplementary Results 2: IFJ to lPFC connectivity

*Objective.* Based on previous findings in age differences between children and adults, where we showed that children showed a greater increase in connectivity between the IFJ and the lateral PFC from single to repeat blocks (Schwarze et al., 2023), we tested whether this connectivity changed with practice.

*Methods*. For our analyses to be less dependent on the precise location of the lPFC cluster at the first timepoint, we defined two anatomical ROIs (in the frontal polar cortex [FPC] and the middle frontal gyrus [MFG]) based on the Harvard-Oxford Atlas (Makris et al., 2006) thresholded at a probability of 30%. In our previous analyses, we had used these two anatomical regions to restrict the cluster that showed greater connectivity with the IFJ in children compared to adults. Thus, the analyses retained the general location of the cluster without being too dependent on the precise location. We extracted PPI parameters for these ROIs from the session-specific repeat > single PPI model described in the main analyses and subsequently analyzed these for changes between sessions A and D and for differences between the SI, the SW, and the PC groups. Models included random intercepts for participants and random slopes for sessions.

*Results*. Neither the IFJ to FPC, nor the IFJ to MFG connectivity showed changes with practice. However, connectivity for repeat trials was greater than for single trials across sessions and groups (FPC: est. = 1.48; 95%-CI: 0.19; 2.74; MFG: est. = 2.26; 95%-CI: 1.04; 3.50), suggesting that also after practice children involved these additional regions with increased mixing demand on repeat trials (see Table SR2.1).

Table SR2.1 Model outputs of PPI parameters across sessions A and D, the three groups (SI vs. PC, SW vs. PC), and repeat and single trials. CI denotes 95%-Credible Intervals; bold values indicate estimates whose 95%-CI did not include zero.

| Effect | IFJ – FPC | | | IFJ – MFG | | |
|---|---|---|---|---|---|---|
| | Estimate | CI | | Estimate | CI | |
| Intercept | -1.73 | -2.86 | -0.61 | -1.84 | -2.94 | -0.73 |
| group: SI vs. PC | 0.33 | -1.16 | 1.84 | 0.38 | -1.11 | 1.87 |
| group: SW vs. PC | 1.17 | -0.31 | 2.63 | **1.67** | 0.22 | 3.14 |
| session: D vs. A | 0.91 | -0.49 | 2.32 | 1.08 | -0.24 | 2.41 |
| condition: repeat vs. single | **1.48** | 0.19 | 2.74 | **2.26** | 1.04 | 3.5 |
| groupSI x sessionD | -0.07 | -1.98 | 1.82 | -1.04 | -2.85 | 0.78 |
| groupSW x sessionD | -0.59 | -2.46 | 1.27 | -1.46 | -3.3 | 0.31 |
| group SI x repeat | 0.4 | -1.35 | 2.1 | -0.56 | -2.21 | 1.13 |
| group SW x repeat | -0.25 | -1.94 | 1.42 | -1.02 | -2.59 | 0.57 |
| session D x repeat | -0.33 | -2.19 | 1.48 | -1.18 | -2.93 | 0.57 |
| group SI x session D x repeat | -0.31 | -2.82 | 2.24 | 0.87 | -1.57 | 3.25 |
| group SW x session D x repeat | -0.33 | -2.77 | 2.11 | 0.47 | -1.83 | 2.79 |

**Supplementary Results 3: Changes in deviation score and association with changes in connectivity**

*Hypotheses*: For detailed hypotheses see pre-registrations at https://osf.io/by4zq/. In brief, based on results of the pre-training session (Schwarze et al., 2023), we expected changes in IFJ–lPFC connectivity to depend on how adult-like the pre-training activation of a child was (Hypothesis 5). Additionally, we expected the alignment of changes in activation pattern and connectivity to be associated with practice-related changes in performance (Hypothesis 6).

*Methods*:
For detailed methods, see pre-registrations at https://osf.io/by4zq/.
*Deviation Score.* Briefly, we computed functional activation deviation scores for repeat > single and switch > repeat activation based on Düzel et al. (2011; see also Fandakova et al. 2015) for each child relative to the adult average of the respective contrast. A more negative score represents more adult-like activation patterns, and a more positive score represents less adult-like patterns.
*Deviation score and change in connectivity*. To test whether change in IFJ–lPFC connectivity depended on how adult-like a child's repeat > single activation at session A was, we performed Bayesian linear mixed modeling with IFJ–lPFC connectivity as the dependent variable, and condition (single vs. repeat), session (A vs. D), and deviation score at the first session (high vs. low, based on a median split) as fixed effects, allowing for all interactions. We analyzed only the two training groups (i.e., SW and SI). The model further included a random intercept of individuals and random slope of session.
Given that neither deviation scores nor IFJ–lPFC connectivity changed with practice, we did not further analyze the preregistered Hypothesis 6.

*Results*: Deviation scores did not change over the course of practice, neither for the repeat > single contrast (Figure SR3.1A), nor the switch repeat contrast (Figure SR3.1B; all 95%-CI included zero). Model outputs are shown in Table SR3.1 for comparisons of session A and D among the three groups (SI vs. PC and SW vs. PC) and in Table SR3.2 for comparisons across all sessions (A, B, C, and D) for the two practice groups (SW vs. SI). The change in connectivity between sessions A and D did not differ between children who showed more and those who showed less adult-like activation patterns at session A (see Table SR3.3).

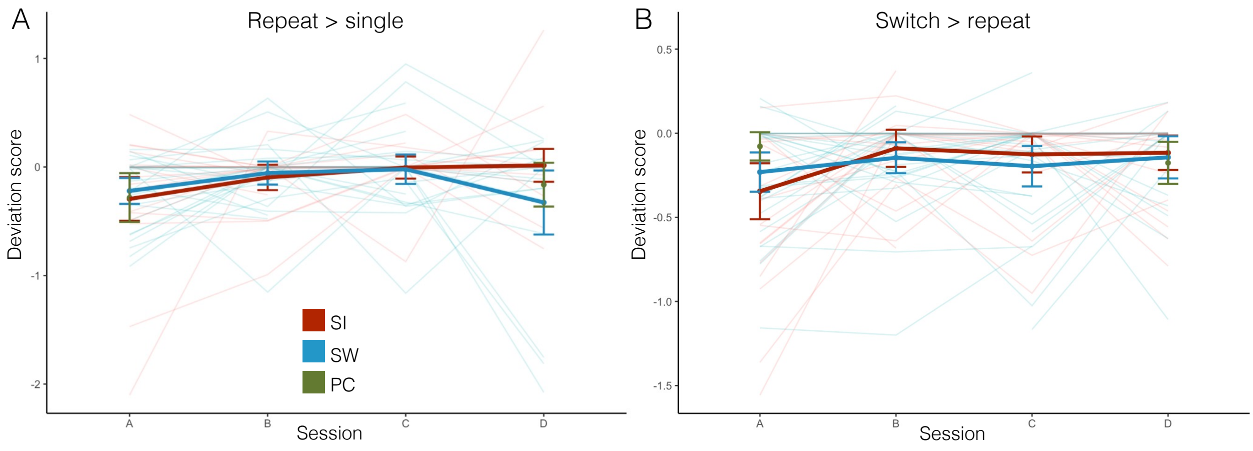

*Figure SR3.1 Changes in deviation score.* (A) Change in deviation score for the repeat > single contrast. (B) Change in deviation score for the switch > repeat contrast. Higher values denote less adult-like activation pattern. Thin lines show individual trajectories. The passive control (PC) group is shown in green, the single-task practice (SI) group in red, and the task-switching practice (SW) group in blue. Error bars denote 95%-confidence intervals.

Table SR3.1. Model output for comparisons of session A and D among the three groups (SI vs. PC and SW vs. PC). CI denotes 95%-Credible Intervals

| Effect | Repeat > Single | | | Switch > Repeat | | |
|---|---|---|---|---|---|---|
| | Estimate | CI | | Estimate | CI | |
| Intercept | -0.01 | -0.04 | 0.02 | -0.01 | -0.10 | 0.02 |
| session: D vs. A | -0.01 | -0.05 | 0.04 | 0 | -0.06 | 0.06 |
| group: SI vs. PC | -0.01 | -0.07 | 0.03 | -0.03 | -0.22 | 0.01 |
| group: SW vs. PC | 0 | -0.05 | 0.04 | -0.01 | -0.09 | 0.03 |
| group SI x session D | 0.02 | -0.04 | 0.10 | 0.03 | -0.03 | 0.23 |
| group SW x session D | 0.01 | -0.05 | 0.08 | 0 | -0.06 | 0.11 |

Table SR3.2. Model output for comparisons across all sessions (A, B, C, and D) for the two practice groups (SW vs. SI). CI denotes 95%-Credible Intervals

| Effect | Repeat > Single | | | Switch > Repeat | | |
|---|---|---|---|---|---|---|
| | Estimate | CI | | Estimate | CI | |
| Intercept | -0.01 | -0.05 | 0.01 | -0.03 | -0.07 | -0.00 |
| session B | 0.01 | -0.03 | 0.05 | 0.02 | -0.04 | 0.08 |
| session C | 0.01 | -0.02 | 0.06 | 0.02 | -0.03 | 0.07 |
| session D | 0.01 | -0.03 | 0.06 | 0.03 | -0.02 | 0.08 |
| group SW vs. SI | 0.01 | -0.03 | 0.05 | 0.02 | -0.02 | 0.07 |
| session B x group SW | 0 | -0.06 | 0.04 | -0.05 | -0.13 | 0.01 |
| session C x group SW | -0.01 | -0.07 | 0.04 | -0.03 | -0.09 | 0.03 |
| session D x group SW | -0.01 | -0.07 | 0.05 | -0.03 | -0.09 | 0.03 |

Table SR3.3. Model output for connectivity at session A and D predicted by deviation score (median split) at session A across both practice groups. CI denotes 95%-Credible Intervals; bold values indicate estimates whose 95%-CI did not include zero.

| Effect | IFJ – FPC | | | IFJ – MFG | | |
|---|---|---|---|---|---|---|
| | Estimate | CI | | Estimate | CI | |
| Intercept | -0.52 | -1.54 | 0.5 | 0.08 | -1.01 | 1.18 |
| session: D vs. A | -0.12 | -1.55 | 1.28 | -1.19 | -2.61 | 0.21 |
| condition: repeat vs. single | 1.13 | -0.02 | 2.29 | 0.52 | -0.63 | 1.67 |
| deviation score group: more vs. less adult-like | -0.78 | -2.18 | 0.6 | **-1.46** | -2.94 | -0.02 |
| session D x repeat | 0.13 | -1.7 | 2 | 0.83 | -0.98 | 2.69 |
| session D x group more adult-like | 1.47 | -0.44 | 3.39 | 1.86 | -0.05 | 3.78 |
| group more adult-like x repeat | 0.77 | -0.78 | 2.34 | **1.7** | 0.16 | 3.25 |
| session D x repeat x group more adult-like | -1.45 | -3.96 | 1.03 | -2.39 | -4.95 | 0.12 |
